# RNA Editors Sculpt the Transcriptome During Terminal Erythropoiesis

**DOI:** 10.1101/2025.04.03.647020

**Authors:** Areum Han, Alena Yermalovich, Mohamad Ali T. Najia, Daniel S. Pearson, Yuko Fujiwara, Michael Bolgov, Caroline Kubaczka, Trista E. North, Vanessa Lundin, Stuart Orkin, George Q. Daley

## Abstract

Selective RNA degradation during terminal erythropoiesis results in a globin-rich transcriptome in mature erythrocytes, but the specific RNA decay pathways remain unknown. We found that deficiency of the terminal uridylyl transferase enzyme Zcchc6 and the 3’-5’ exoribonuclease Dis3l2 in mouse models led to fetal and perinatal reticulocytosis, an accumulation of RNA-rich precursors of terminal erythroid cells, suggesting their crucial roles in terminal red cell differentiation. Notably, knockout embryos exhibited persistent high-level expression of *Hbb-bh1* globin, the ortholog of human fetal *γ-*globin. Perturbation of the Zcchc6-Dis3l2 pathway in mice engineered to express the human β-globin locus likewise increased *γ*-globin levels in fetal erythroid cells, suggesting that globin switching entails post-transcriptional mechanisms of mRNA destabilization in addition to transcriptional down-regulation. We cultured human hematopoietic stem and progenitor cells (HSPCs), performed CRISPR/Cas9-mediated knockout of ZCCHC6 and DIS3L2, and observed accumulation of RNA and elevated γ-globin levels in terminal erythroid cells. Our findings reveal a conserved role for the ZCCHC6/DIS3L2 RNA editors in terminal erythropoiesis and demonstrate a post-transcriptional mechanism for *γ-*globin gene switching, advancing research into *in vitro* erythrocyte generation and *γ-*globin stabilization to ameliorate hemoglobinopathies.

## INTRODUCTION

Terminal differentiation of red blood cells (RBCs)^1,2^ entails the loss of key cellular components such as the nucleus, mitochondria, and ribosomes^3–7^. After enucleation, orthochromatic erythroid cells become reticulocytes, which then undergo extensive RNA degradation to develop into mature erythrocytes, primarily composed of globins^8^. Previous examination of RNA half-life suggests that active decay of non-erythrocyte transcripts contributes to this globin-rich, low-entropy transcriptome^9,10^, in addition to global proteome remodeling through protein tagging by ubiquitination^11^.

The Terminal Uridylyl Transferase ZCCHC6 (TUTase 7) has recently been implicated in regulating the degradation of various RNAs during cellular transitions, rather than in steady-state conditions^12–20^. This re-sculpting of the transcriptome occurs in transcriptionally compromised or quiescent contexts, such as when cells are treated with the transcription inhibitor Actinomycin D^21^ or during the maternal-to-zygotic transition (MZT) ^22^. During MZT, a sharp decline in maternal transcripts accelerates the emergence of the zygotic transcriptome. This coordinated transcriptome remodeling is comparable to the terminal stages of erythropoiesis.

Embryonic, fetal, and adult hemoglobins are sequentially expressed during mouse and human erythroid ontogeny ^23^. In humans, during the switch from fetal globin (HbF; α2γ2) to adult globin (α2β2), γ-globins are replaced by adult β-globins. Conversely, faulty adult β-globin with pathogenic mutations can be rescued by reactivating γ-globin expression. Indeed, elevated γ-globin levels in adults have been shown to ameliorate symptoms of sickle cell disease ^24,25^. Due to this therapeutic potential, the transcriptional regulation of γ-globin has been extensively studied ^26,27^, while its post-transcriptional regulation remains less explored ^28,29^.

Prior studies have shown that the ZCCHC6 adds uridines to the 3’ end of target RNAs, which serves as a recognition signal for the 3′ to 5′ exonuclease DIS3L2, leading to the degradation of target RNAs ^12,18,30,31^. *Zcchc6*, uniquely recognized for its RNA-tagging capability, is upregulated in the peripheral blood of mice exposed to chronic hypoxia and subsequently downregulated during normoxic recovery ^32^, suggesting that increased levels of ZCCHC6 might be necessary to process the higher number of reticulocytes generated during erythropoietic stress. ZCCHC6 appears to be a rate-limiting factor in RNA editing, as it is slower than nuclease activity. Based on all the above, we posited that the ZCCHC6-DIS3L2 axis might regulate RNA fate during terminal erythropoiesis, accelerating RNA degradation and potentially influencing globin switching. In this study, we explored the role of the RNA editor-exonuclease axis during terminal erythropoiesis using murine models and in vitro erythroid differentiation of human blood progenitors. Our findings revealed that loss of this axis leads to RNA accumulation and upregulation of fetal globin expression, resulting in a reversed globin switching pattern during terminal erythropoiesis. This highlights a novel, conserved post-transcriptional gene regulation pathway during late erythropoiesis.

## RESULTS

### Whole-body *Zcchc6/Dis3l2* KO mice exhibit reticulocytosis and anemia

We began our investigation by analyzing the expression profiles of Zcchc6 and Dis3L2 throughout erythropoietic ontogeny, examining blood from knockout (KO) models across embryonic stages to adulthood. Analysis of a public dataset demonstrated that the expression of *Zcchc6* is upregulated as erythroid cells differentiate into the terminal stages of fetal and adult definitive erythropoiesis^33,34^ (Extended Data Fig. 1, upper panel), suggesting that Zcchc6 plays an important role in terminal erythropoiesis. In contrast, *Dis3l2* did not show high expression in the terminal stages (Extended Data Fig. 1, lower panel). As an exonuclease, Dis3L2 may not be a limiting factor in RNA regulation.

We generated a whole-body *Dis3l2* KO model (Extended Data Fig. 2), but the *Dis3l2* KOs exhibited postnatal lethality. Adult *Zcchc6* KOs survived and demonstrated an increase in reticulocytes compared to wild-type (WT) controls, suggesting a role for Zcchc6 during the transition from reticulocytes to mature RBCs (Extended Data Fig. 3c). Other hematological parameters in the *Zcchc6* KO were mostly comparable to those of the WT in young adult stages (Extended Data Fig. 3 a&b). Zcchc6 KO animals exhibited increased neutrophil counts compared to heterozygous (HET) animals (Extended Data Fig. 3b).

At embryonic day (E) 18.5, cells are undergoing RNA degradation as they differentiate from reticulocytes to erythrocytes, and both KOs are viable. Therefore, we investigated animals at this age and observed embryonic anemia and reticulocytosis in both KO models. In detail, we collected peripheral blood and spleen cells from E18.5 embryos, and performed histology, flow cytometry, and hemogram analysis (Fig.1a).

Examination of blood smears from *Zcchc6* and *Dis3l2* KO embryos showed polychromasia and increased RNA retention in red cells, as revealed by staining with methylene blue, indicating reticulocytosis (Fig.1b). We quantified reticulocytosis using flow cytometry, pre-gating erythroid (TER119+) cells and staining for the CD71 transferrin receptor and the nucleic acid dye Thiazole Orange (TO) ^35,36^ (Fig.1c). WTs displayed expected patterns of CD71 loss and RNA (TO signal) loss, while KOs showed higher levels of RNA and a significantly higher CD71^low^TO^high^ population relative to HETs and WTs (Fig.1c). These data demonstrate that Zcchc6 and Dis3l2 impact RNA degradation during terminal erythropoiesis. We also profiled erythroid RNA dynamics in the embryonic spleen, an erythropoietic organ at E18.5^37^. Embryonic spleen harbors nucleated erythroid cells, and we observed a large drop in TO signal in the CD71^high^ population due to enucleation^35^ (Extended Data Fig.4a). However, we did not observe differences in enucleation ratio across genotypes. On the other hand, in the later stage of erythroid cell differentiation (TER119^+^CD71^low^), the CD71^low^TO^high^ population was elevated in KOs compared to HETs and WTs (Extended Data Fig.4a), again indicating inefficient RNA degradation in KOs. In both peripheral blood and spleen, this accumulation of the CD71^low^TO^high^ population was genotype-dependent on Zcchc6 (Extended Data Fig.4a, right graph). This result suggests that cellular levels of Zcchc6 determine the extent of RNA degradation.

Both KOs at E18.5 had reduced RBC numbers, decreased hematocrit (HCT) and increased Mean Corpuscular Hemoglobin (MCH) compared to HETs and WTs (Fig.1d). *Zcchc6* KO also showed increased Red Cell Distribution Width (RDW); a trend observed in *Dis3l2* KO that failed to reach statistical significance in WT. For *Dis3l2* KOs, we observed a significant increase in Mean Corpuscular Volume (MCV) and lower hemoglobin concentration (Hb), a trend also observed in *Zcchc6* KOs but without statistical significance. Both KOs were anemic from a reduced number of RBCs, and both showed lymphocytosis (Extended Data Fig. 4b). *Zcchc6* KO generally showed more marked differences in the leukocyte panel with an increased absolute number and percentage of monocytes (MO), a decreased percentage of basophils (BA), and increased number in neutrophils (NE). These interesting observations could be further examined in future studies.

### Conditional *Zcchc6/Dis3l2* KO mice phenocopy whole-body KOs

To examine erythrocyte-specific phenotypes, we utilized animals in which the *Zcchc6/Dis3l2* alleles were flanked by LoxP sites (floxed) to generate conditional KOs of these genes by crossing to strains expressing *EpoR-Cre* ^38^ (Fig. 2a & Extended Data Fig. 2). Conditional KO in erythroid cells (TER119+) was confirmed by western blot (Extended Data Fig. 2b). Indeed, cKOs phenocopied the reticulocytosis observed in whole-body KOs (Fig. 2b), indicating that this phenotype is cell-autonomous. Both cKOs exhibited a statistically significant increase in the CD71^low^TO^high^ population in peripheral blood (Fig. 2b) and spleen (Extended Data Fig. 5a) compared to *EpoR-Cre*-carrying and cWT animals. Hematological readings in both cKOs showed consistent reduction in RBC numbers (Fig. 2c). cKOs again showed a trend of increased absolute WBC numbers, specifically in the absolute number of lymphocytes (Extended Data Fig. 5b).

**Fig. 1.**
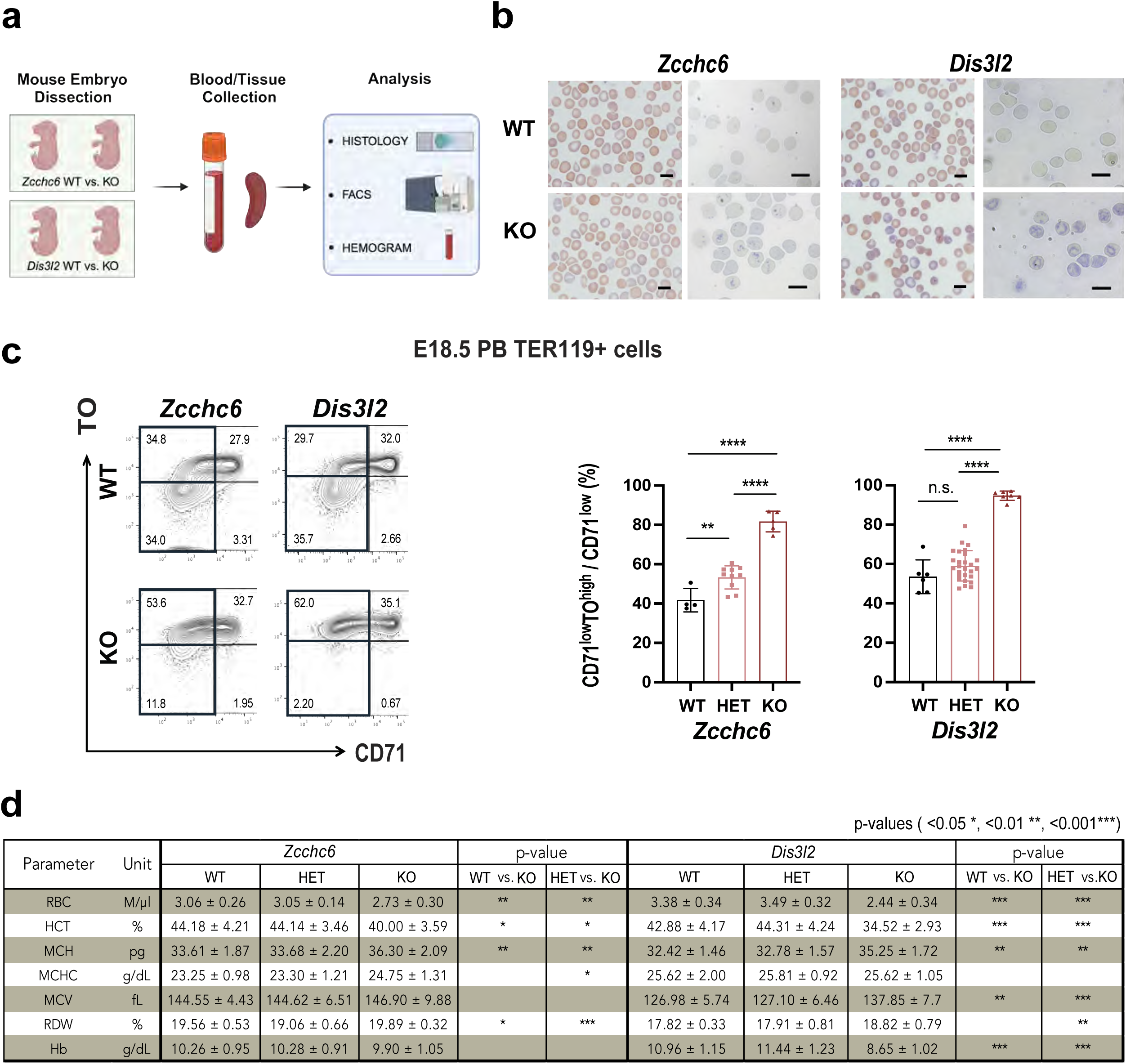
Analysis of Whole-Body *Zcchc6/Dis3l2* Knockout Mice. **(a)** The overall scheme of the embryonic blood studies on *Zcchc6* and *Dis3l2* knockout (KO) mice. **(b)** Histology of peripheral blood smears stained by May-Grünwald-Giemsa (left) and methylene blue (right). Scale bar = *10um*. **(c, left figure)** Flow cytometry profiles of erythroid cells from peripheral blood (PB). Representative profiles of each KO and littermate WT were displayed. **(c, right graphs)** Quantification of reticulocyte population per genotype (mean ± SD). Multiple litters were used. **(d)** Hematological profiles of the peripheral blood (mean ± SD). HCT; Hematocrit, MCH; Mean corpuscular hemoglobin, MCHC; Mean corpuscular hemoglobin concentration, MCV; Mean corpuscular volume, RDW; Red cell Distribution Width.

**Fig. 2.**
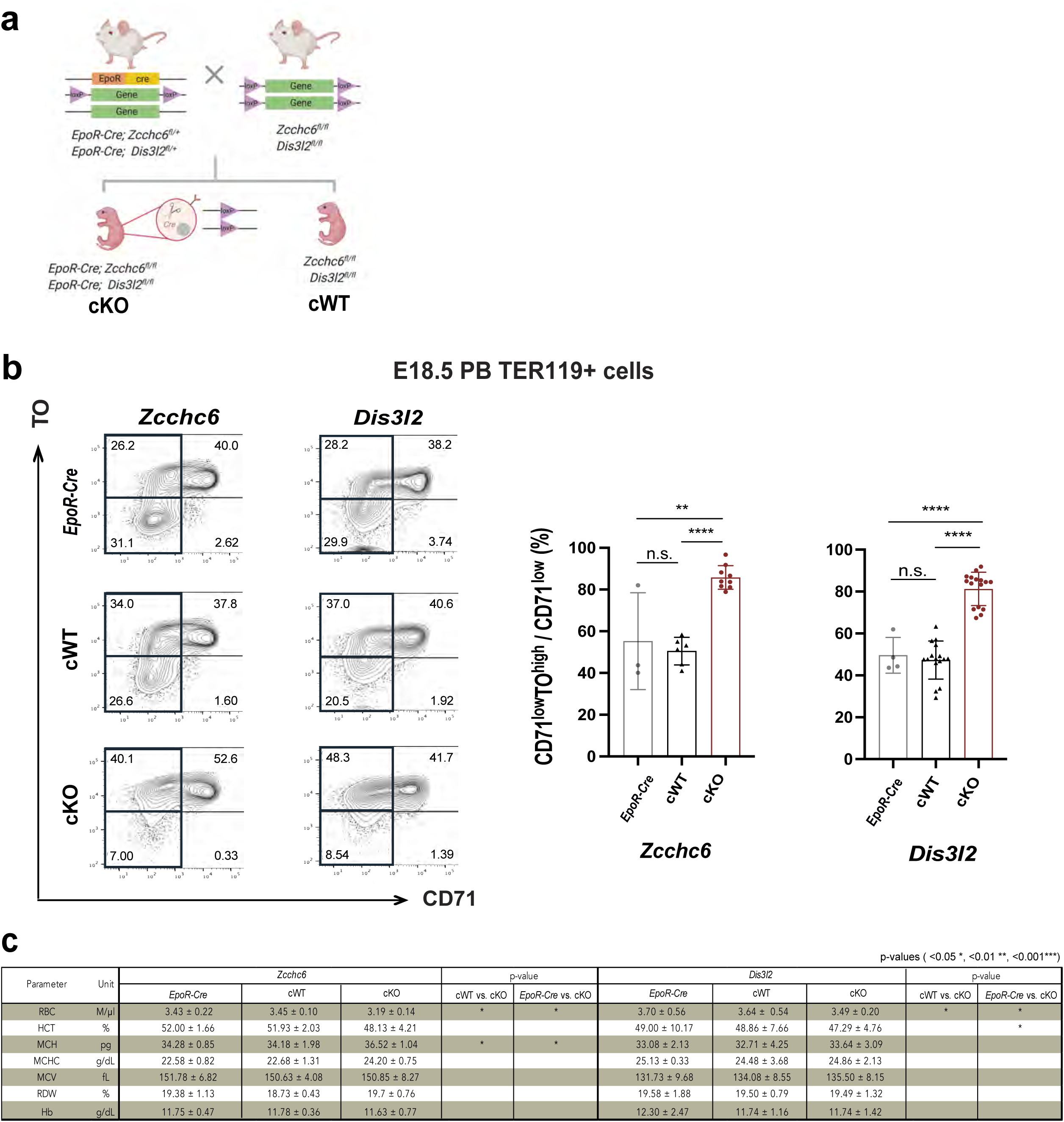
Analysis of Conditional *Zcchc6/Dis3l2* Knockout Mice. **(a)** Breeding scheme to generate erythroid-specific, conditional KO mice. **(b, left figures)** Representative FACS profiles from peripheral blood of embryos carrying only the *EpoR-Cre* allele (top lane), only the flox allele (cWT, middle lane), and both alleles; erythroid-specific conditional KOs (cKO, bottom lane). **(b, right graphs)** Quantification of reticulocyte (CD71^low^TO^high^) population per genotype. **(c)** Hematological profile results from the PB of conditional KOs (erythrocyte panel).

To demonstrate that the observed reticulocytosis reflected failed red cell maturation and not hemolytic anemia, we confirmed preserved levels of haptoglobin and the absence of elevated LDH and bilirubin levels ^39^ (Extended Data Fig. 6). Taken together, these results indicate that the observed reticulocytosis arises from an RNA degradation defect intrinsic to the erythroid cell during terminal differentiation.

### Normal terminal erythropoiesis in KO mice lacking *Lin28a/b* and *Zcchc11*

An enzymatic axis regulating microRNA biogenesis has been described whereby the RNA binding proteins LIN28A/B dock to the loop of the microRNA precursor *pre-let-7*, which then recruits ZCCHC6 or the paralog ZCCHC11, resulting in an inhibitory tri-complex^18^. The assembled tri-complex facilitates oligouridylation, and the oligouridylated RNA species then become substrates for DIS3L2^30^ (Extended Data Fig. 7a). However, neither *Lin28a* nor *Lin28b* KOs exhibited reticulocytosis (Extended Data Fig. 7b). Paralogs often perform similar functions and manifest redundancy. Among seven TUTases in humans, ZCCHC6 and ZCCHC11 are the two largest TUTases with CCHC and C2H2-type zinc finger domains^18^. Previous studies have shown that ZCCHC6 and ZCCHC11 have redundant RNA editing activity ^21,31,40^. Importantly, while *Zcchc11* heterozygotes exhibited slight reticulocytosis, this was not observed in *Zcchc11* KOs, suggesting that Zcchc11 may provide at best partial redundancy (Extended Data Fig. 7b). The *Zcchc6* and *Zcchc11* quad-flox system also showed that significant reticulocytosis was observed only in animals with *Zcchc6* cKO (homozygous floxed alleles of *Zcchc6* driven by *EpoR-Cre)*, but not in *Zcchc11* cKO (Extended Data Fig. 8).

### Reticulocytosis showed postnatal attenuation

*Dis3l2* whole-body KO mice die shortly after birth, whereas *Dis3l2* cKOs under *EpoR-Cre* survive postnatally, allowing us to investigate adult peripheral blood. Both *Zcchc6 KO* and *Dis3l2* cKOs showed significant reticulocytosis at Postnatal day (P) 2 and P21 (Extended Data Fig. 9). Interestingly, the degree of reticulocytosis was attenuated with age. We postulate there may be differential RNA regulation in the bone marrow and adult peripheral blood compared to the embryonic spleen and blood.

### Characterization of accumulated RNA in *Zcchc6* and *Dis3l2* KO mice

Next, we investigated the nature of RNA accumulated in *Zcchc6* and *Dis3l2* KOs at E18.5 utilizing RNA-seq (Fig. 3). The estimated expression levels in *Zcchc6* and *Dis3l2* KOs clustered by KO genotype (Extended Data Fig. 10b). In PCA analysis, the *Dis3l2* KO differed more from the WT than did the *Zcchc6* KO (Fig. 3b), consistent with more severe RNA accumulation in *Dis3l2* KO than in *Zcchc6* KO. Differential gene expression analysis also showed that *Dis3l2* KOs had more differentially expressed transcripts than did the *Zcchc6* KOs (Fig. 3c). Both *Zcchc6* and *Dis3l2* KOs had more over-represented transcripts (3.0% and 9.6%) than under-represented transcripts (1.3% and 5.3%). Downregulated transcripts lacked any significant gene ontology categories, suggesting a lack of target specificity for the population of affected mRNAs. On the other hand, upregulated genes for each KO demonstrated some gene ontology term enrichment. “Cytoplasmic translation” and “ribonucleoprotein complex biogenesis” were the most enriched terms in *Zcchc6* KO, whereas enriched terms for *Dis3l2* KO were “regulation of the mRNA metabolic process”, “proteasomal protein catabolic process”, “NIK/NF-kappaB signaling”, “generation of precursor metabolites and energy”, and “ribonucleoprotein complex subunit organization” (Extended Data Fig. 10c). When the differentially expressed genes from both KOs were overlaid (Fig. 3d), we found that most genes (75% of the total) were aligned in the same regulatory direction, with the majority being accumulated in both KOs. This included the non-coding RNA *H19*, a known target of Dis3l2, identified in a previous nephron progenitor study ^41^. “Cytoplasmic translation” and “ribonucleoprotein complex regulation” were gene ontology terms common to both KOs (Fig. 3e). We further validated the top upregulated and downregulated genes by performing qPCRs on cKO samples (Fig. 3f). The direction of regulation was consistent, with several genes significantly upregulated in both cKOs: *Hbb-bh1*, *Ccnd3*, *Gnb2*, *Hspa8*, and *Alad*. The *Hbb-bh1* transcript was more upregulated in the *Dis3l2* KO than in the *Zcchc6* KO, consistent with our observation of greater RNA accumulation in the *Dis3l2* KO. *Hbb-bh1* is the embryonic/fetal globin, which is unusual to observe at our dissection time point of E18.5, as globin switching has already occurred in wild type mice by this point ^42,43^.

**Fig. 3.**
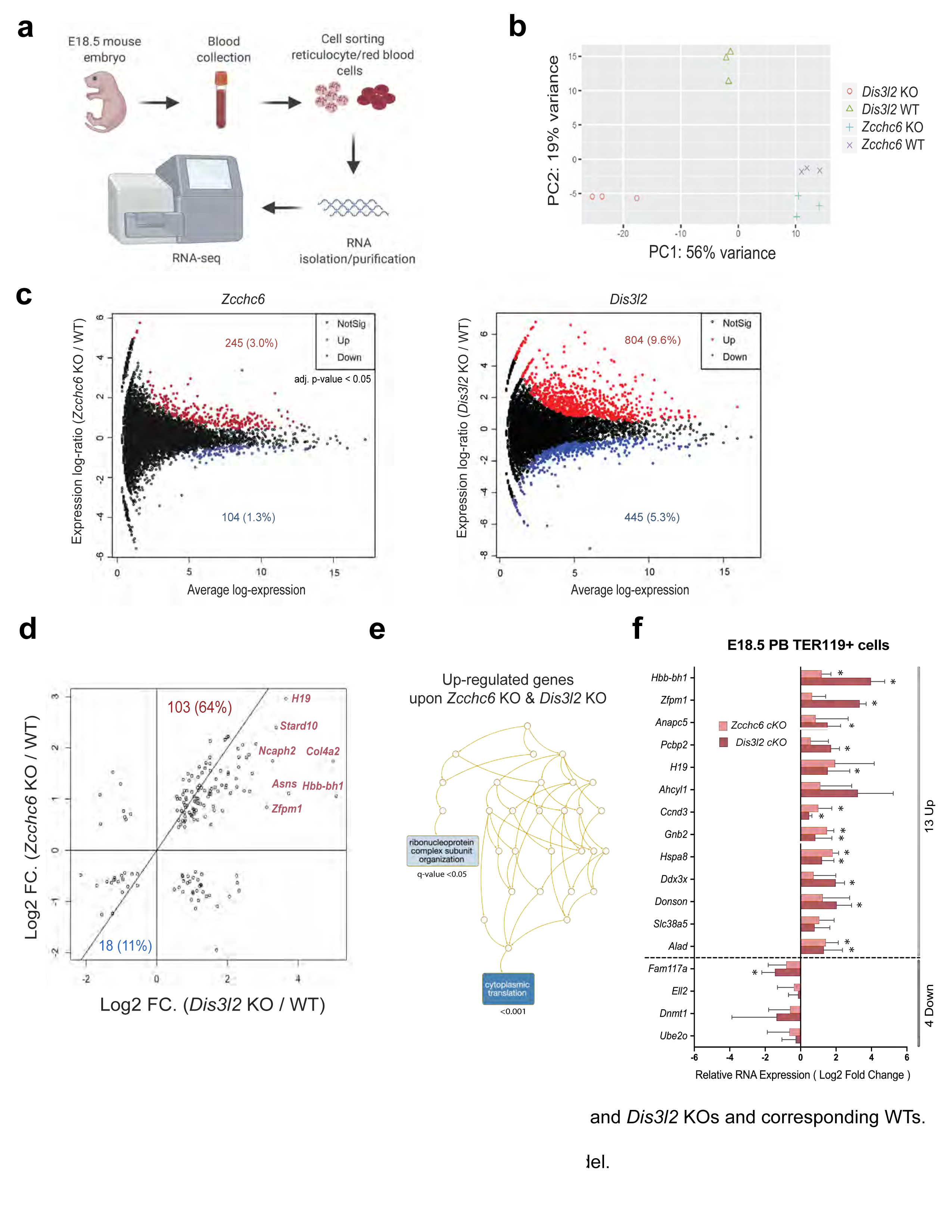
RNA Profiling. **(a)** RNA-seq experimental scheme from embryonic erythroid cells. **(b)** Principal component analysis of global transcriptome from *Zcchc6* and *Dis3l2* KOs and corresponding WTs. **(c)** Differentially expressed genes in each KO model. **(d)** Overlap analysis of differentially expressed genes in each KO model. **(e)** Gene ontology analysis of up-regulated genes from both KOs. **(f)** Validation of top hits using qPCRs with samples from cKOs.

### Upregulation of *γ-*globin in *Zcchc6* and *Dis3l2* KOs in Humanized Mice

Hbb-bh1 is the ortholog of the human fetal γ-globin (gamma; *HBG1/HBG2*). Research has established that patients with pathogenic sickle cell mutations in adult β-globin show improved symptomology following reactivation of fetal γ-globin^25,44^. Thus, we aimed to test whether *Zcchc6/Dis3l2* deficiency would upregulate the human γ-globin gene. We crossed our *Zcchc6* and *Dis3l2* KOs with a humanized β-YAC model mouse harboring the human β-globin gene cluster as a yeast artificial chromosome transgene ^45^ (Fig. 4a) and validated the new model through genomic and protein analysis (Extended Data Fig.11a, b). The human *γ*-globin was significantly upregulated in erythroid cells at E18.5 in the *Zcchc6* and *Dis3l2* KOs (Fig. 4b). These humanized β-YAC mice also preserved human globin switching in the mouse, enabling the profiling of the dynamics of all β-like globins^45^. We examined erythroid cells at an earlier time point (E14.5 fetal liver) and observed up-regulation of human fetal *γ*-globins and down-regulation of adult *β*-globin relative to WT (Fig. 4c). At E18.5, this pattern of reverse switching persisted in the peripheral blood, albeit to a lesser degree (Fig. 4d), and was independent of maturation differences between KO and WT mice (Extended Data Fig. 11 c&d). Together, these results highlight that ZCCHC6/DIS3L2 upregulates human fetal γ-globin in a conserved manner.

**Fig. 4.**
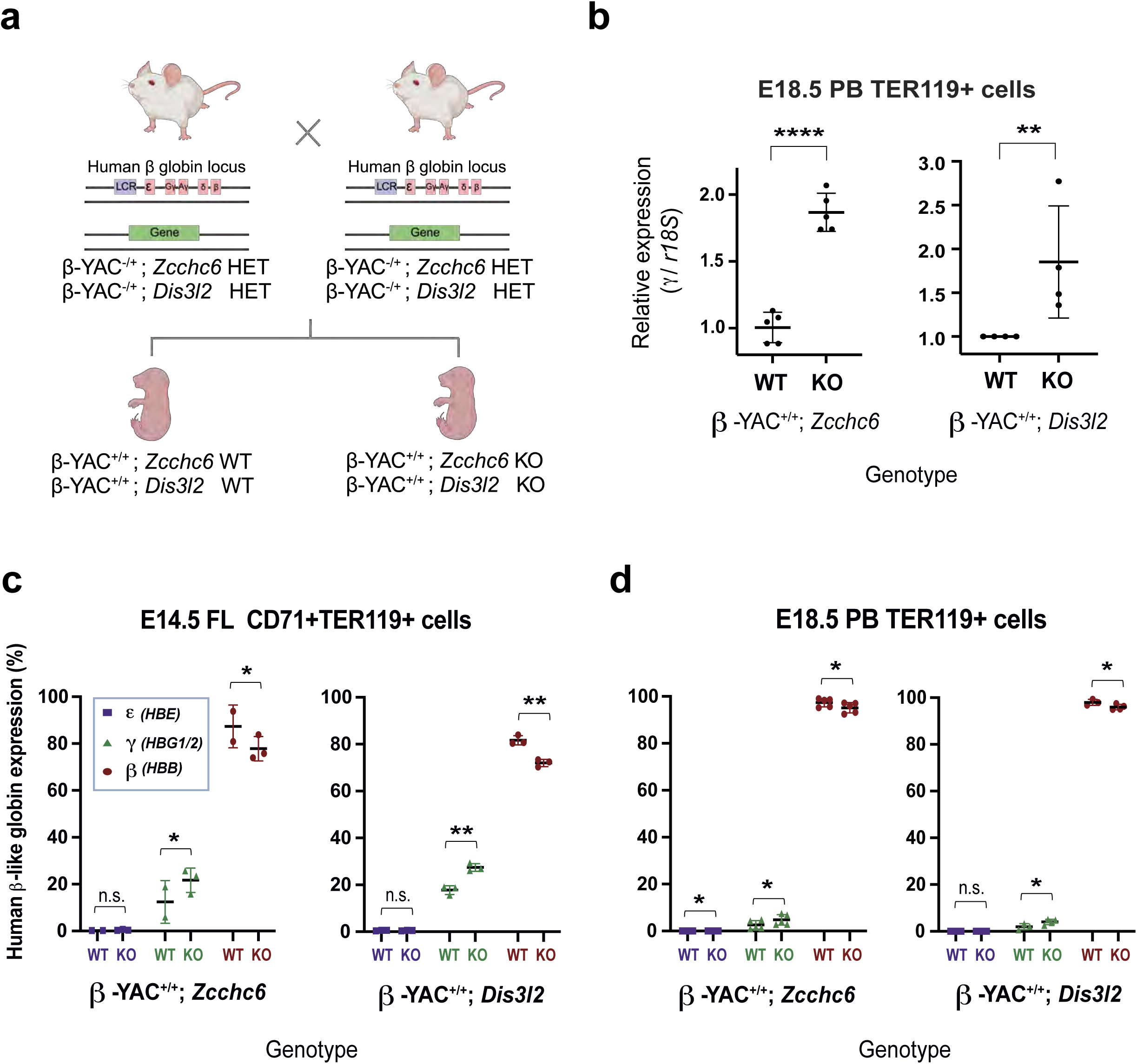
Humanized Murine Model Study. **(a)** Breeding scheme for crossing *Zcchc6/Dis3l2* KO mice with humanized globin YAC mice. **(b)** Fold change in normalized *γ-globin* expression (Ct values for *γ-globin* transcripts were first normalized to those of *r18S* transcripts, and the fold change was then calculated relative to the value of littermate WTs). **(c-d)** Human *β-like* globin profiles on sorted erythroid cells from E14.5 fetal liver and E18.5 PB of humanized mouse embryos using qPCRs. For each embryonic age and mouse model, embryos from two litters were used.

### Profiling ZCCHC/DIS3L2 during human *in vitro* erythropoiesis

Following our mouse studies, we adopted a human *in vitro* system for erythropoiesis that enables a single, synchronized wave of differentiation, as described^46^, to allow detailed molecular studies. We performed *in vitro* erythropoiesis with human umbilical cord blood-derived CD34+ cells and profiled protein expression and globin mRNA expression (Extended Data Fig. 12). Flow cytometry profiles of differentiating erythroid precursors from cord blood mirrored that of native erythroid ontogeny, with expression of CD71, followed by glycophorin A, and ultimately incomplete terminal loss of CD71 compared to adult peripheral blood (Extended Data Fig. 12b). Immunoblotting showed increasing pan-hemoglobin levels through D20, followed by decreased levels at D23 when some cell lysis occurs. DIS3L2 was highly expressed starting on D9 and thereafter decreased, while ZCCHC6 was highly expressed on D12 and D16, and decreased thereafter (Extended Data Fig. 12c).

Expression of mRNA for HBE (*ε*) and HBG1/2 (*γ*) increased until terminal stages when levels dropped in the setting of cell lysis (Extended Data Fig. 12d). The expression of HBB (β-globin) increased significantly until D16, after which it declined. Histological view at D12 and D16 revealed that erythroid cells become extensively hemoglobinized and exhibit highly condensed or absent nuclei (Extended Data Fig. 12e).

### CRISPR KOs during *in vitro* erythropoiesis phenocopy KO mice

Next, we used CRISPR/Cas9 to generate KOs by introducing ribonucleoprotein complexes targeting ZCCHC6, DIS3L2, and the AAVS1 locus as a control (Fig. 5a) ^47,48^. We conducted genomic analysis at D12 of differentiation, selected samples with over 60% frameshift indels, and confirmed significantly decreased protein levels (Extended Data Fig. 13a&b). KO of the RNA editing machinery did not impact maturation profiles, as reflected by flow cytometry for CD71 and GlyA (Extended Data Fig. 13c). However, we observed significant accumulation of RNA, as reflected by staining with TO (Fig. 5b). In both ZCCHC6 and DIS3L2 KOs, fetal *γ*-globin was upregulated while adult *β*-globin was relatively downregulated (Fig. 5c). However, the effect size was greater for KO of ZCCHC6 relative to DIS3L2. Taken together, these data demonstrate RNA accumulation in CRISPR knockout (KO) cells, which is consistent with the KO mice study, and a reversed globin-switching pattern in these cells, aligning with the humanized murine model study.

**Fig. 5.**
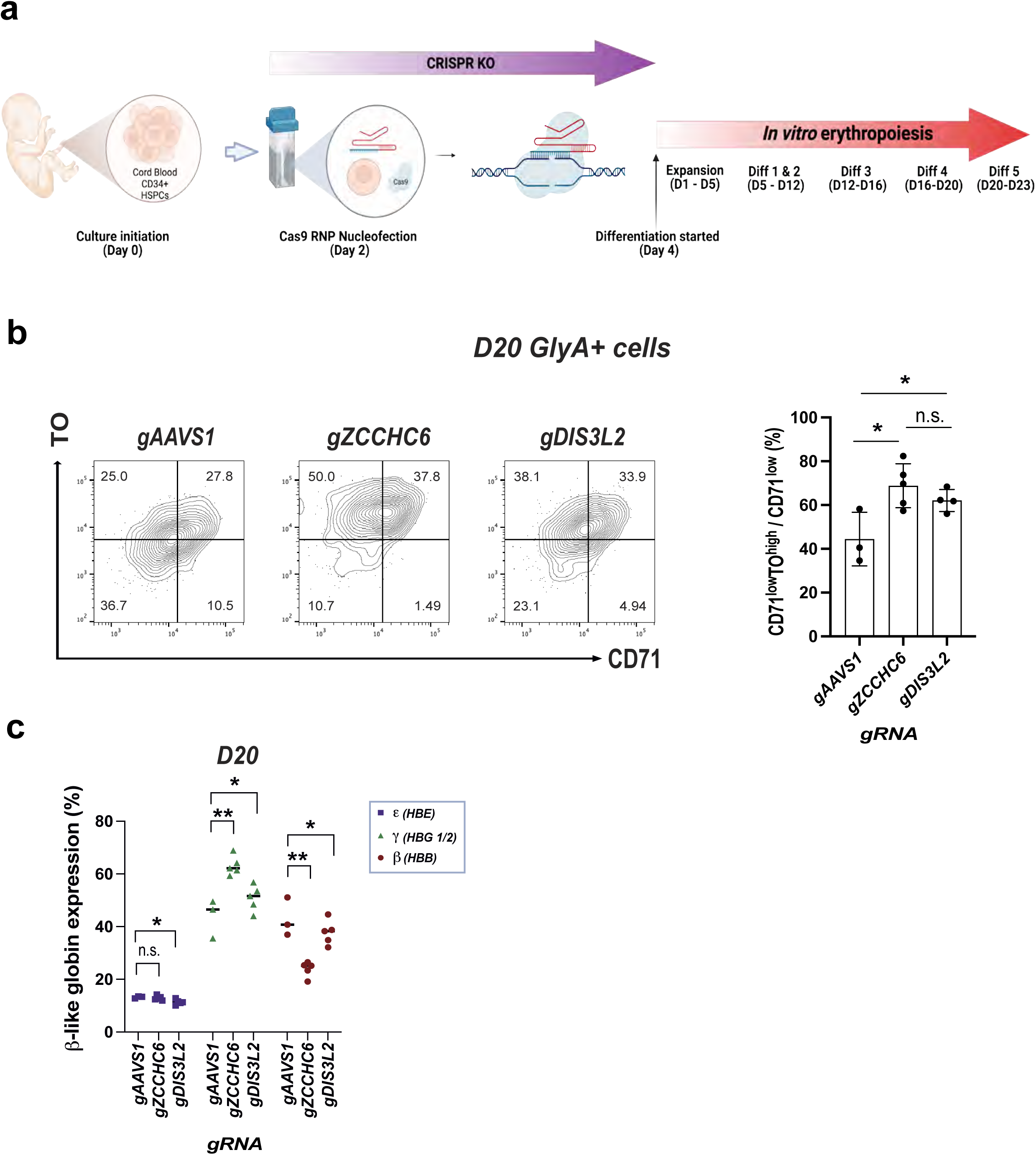
Human *In Vitro* Erythropoiesis Study Using CRISPR Knockout. **(a)** Scheme of the CRISPR/Cas9 knockout followed by *in vitro* erythropoiesis. **(b)** Representative profiles of TO and CD71 within erythroid (GlyA+) cells and the corresponding graph with statistical assessment. **(c)** Globin profiles upon CRISPR/Cas9 knockouts.

### *In vitro* erythroid cells from PB with CRISPR KOs and HU treatment

Reactivation or elevation of the expression of human fetal *γ*-globin by genetic or pharmacologic means is an effective therapeutic strategy in sickle cell disease and transfusion-dependent thalassemia ^49^. Hydroxyurea (HU) treatment is known to elevate γ-globin levels in the peripheral blood of patients with hemoglobinopathies ^50,51^. While a direct mechanistic connection between HU and ZCCHC6 has not been described, a previous study revealed that HU-induced DNA replication arrest triggers the rapid degradation of histone mRNA, mediated by uridylation from ZCCHC6 ^52^.

In this experiment, we aim to test whether the genetic deficiency of ZCCHC6/DIS3L2 affects PB (Peripheral Blood) HPSCs, which better reflect a clinical setting compared to CB HPSCs due to their low baseline levels of fetal hemoglobin. The PB CD34+ cells were subjected to CRISPR knockout (KO) of the RNA editing machinery and differentiated after day 9 in the absence or presence of HU (Fig. 6a) ^53,54^. Consistently, accumulation of reticulocytes (GlyA+CD71^low^TO^high^ cells), elevation of γ-globin levels, and reversed switching patterns were noted in ZCCHC6 CRISPR KOs (Fig. 6b, c, d), while DIS3L2 CRISPR KOs showed a similar trend with a marginal p-value for the first two features. Upon DIS3L2 CRISPR KOs, adult β-globin was also upregulated, resulting in our inability to observe reverse-switching. Genomic analysis at day 12 demonstrated similar frame-shift editing rates in both PB and CB CD34+ cells. No differential impact on erythroid maturation profiles were observed upon CRISPR KOs and HU treatment (Extended Data Fig. 14a & b).

**Fig. 6.**
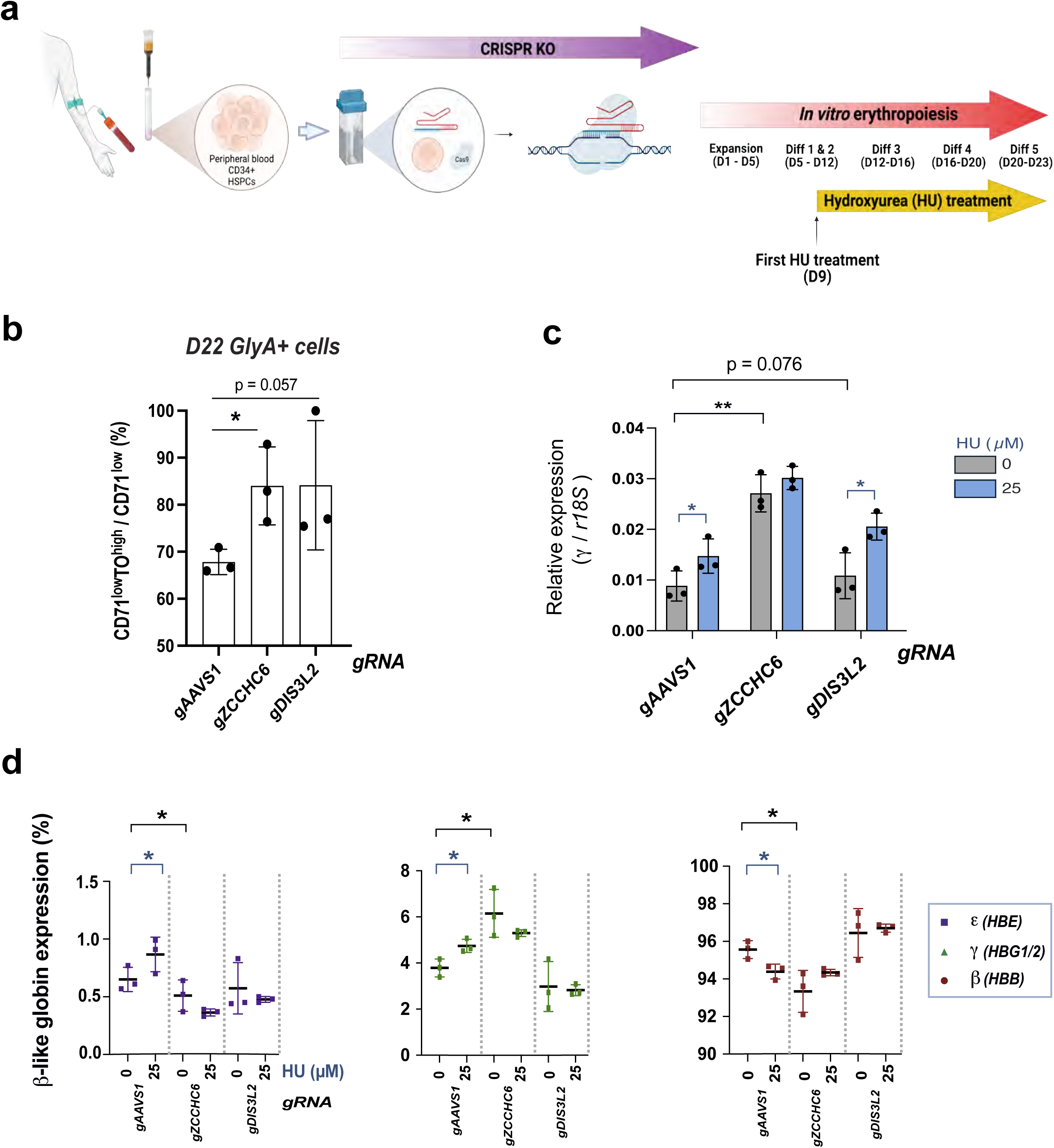
*In Vitro* Erythropoiesis Study with CRISPR KO and HU(Hydroxyurea) Treatment. **(a)** Scheme of the CRISPR/Cas9 knockout and HU treatment followed by *in vitro* erythropoiesis. **(b)** Summary graph and statistical assessment of CD71^low^TO^high^ population upon CRISPR/Cas9 KOs. **(c)** Human *γ-globin* mRNA expression upon CRISPR/Cas9 knockouts and HU treatment. **(d)** Globin profiles upon CRISPR/Cas9 KOs and HU treatment.

When compared with HU treatment, KO of ZCCHC6 alone showed better upregulation of γ-globin levels and reverse switching between γ and β globins than HU treatment (Fig. 6c &d). Furthermore, HU treatment did not augment the impact, likely because the cells reached the maximum capacity to carry fetal globin. Taken together, these results highlight that ZCCHC6 is a consistent regulator in both CB and PB-derived cells. Our data implicating ZCCHC6 as a regulator of γ-globin mRNA stabilization suggests that ZCCHC6 may be a target worth testing further as a therapeutic strategy for enhancing γ-globin expression in hemoglobinopathies.

## DISCUSSION

During terminal differentiation, erythroid cells dramatically discard cellular components, including the nucleus, mitochondria, and ribosomes^3–7^. Nuclear condensation and enucleation entail a loss of transcription^1,3,4^, implicating post-transcriptional regulatory mechanisms as prominent drivers of transcriptome shedding. The resulting transcriptome, with its low-entropy, globin-rich composition in terminal erythroid cells, is thought to be achieved by the active decay of RNAs that are nonessential for mature RBC function^9,10^. However, the exact molecular pathways driving this process have not been elucidated. In this study, we discovered that the ZCCHC6-DIS3L2 RNA editing pathway plays a role in RNA shedding during terminal erythropoiesis, with a prominent target for degradation being fetal (γ-) hemoglobin. Our finding complements the previous discovery of global proteome shedding through ubiquitination^11^, as both mechanisms involve post-translational and post-transcriptional tagging, respectively, to target molecules for degradation. Our finding begs the question of how β-globin mRNA escapes this shedding process. Previous studies have shown that pyrimidine-rich cis-elements in the 3’ UTR, along with associated trans-elements such as αCP and nucleolin, stabilize β-globin mRNAs ^55,56^. This RNP complex may outcompete ZCCHC6 or form an untaggable substrate for the ZCCHC6-DIS3L2 pathway.

Hereditary Persistence of Fetal Hemoglobin (HPFH), a benign genetic condition manifesting as incomplete globin switching, mediates significant amelioration of symptoms when inherited along with the genetic mutation in Sickle Cell Disease (SCD)^25^. This experiment of nature has inspired a therapeutic strategy to reactivate fetal γ-globin via genetic perturbation of the γ-globin repressor BCL11A, resulting in transcriptional reactivation and considerable clinical improvement for SCD patients^44,49,57,58^. Reactivation of γ-globin by genetic editing has achieved proof of concept for treating hemoglobinopathies, marking a seminal translational application of gene editing technology ^44,49^. However, it is challenging to deliver genetic therapy for SCD patients in under-resourced areas of the globe, where SCD is most prevalent ^59,60^. This challenge arises from the scarcity of bone marrow transplantation centers and trained healthcare providers^59,60^, and by the high costs of genetic therapies^61^. The development of small molecule inhibitors that can reactivate γ-globin expression, including inhibition of BCL11A, remains an appealing alternative that could be more readily distributed in under-resourced locales^62–66^. However, achieving therapeutic levels of γ-globin reactivation with a single drug has, to date, proven challenging due to the suboptimal potency of monotherapy and heterogenous pathological symptoms^65^. The successful treatment may therefore require the use of multiple pharmacological agents^65^.

Here, we report a novel post-transcriptional mechanism whereby γ-globin mRNA is subject to active degradation. This discovery raises the possibility that therapeutic strategies to stabilize γ-globin mRNA could synergize with pharmacologic strategies that reactivate γ-globin mRNA expression to ameliorate the symptoms and deleterious health consequences of hemoglobinopathies. Recent studies have reported small molecule activators of γ-globin expression^65,66^, as well as inhibitors of ZCCHC6/11 which have been developed as anti-cancer agents^67,68^. In the latter case, inhibitors reduced the uridylation of target RNAs, including *pre-let-7*^67^, and demonstrated antiproliferative activity in focadhesin (FOCAD)-deleted cancer^68^. Testing existing ZCCHC6 inhibitors or developing novel inhibitors using assays with γ-globin RNA as a substrate in erythroid cells could offer an accessible pharmacological option.

While we observed delayed globin switching, prolonged γ-globin mRNA half-life, reticulocytosis and anemia in the embryonic circulation of *Zcchc6* and *Dis3l2* KO mice, these phenotypes were attenuated in post-weanling mice, when fetal erythropoiesis is fully extinguished in favor of adult patterns of red cell development. Resculpting of the terminal transcriptome during adult murine erythropoiesis most likely entails an additive or distinct regulatory mechanism. In our *in vitro* studies of human erythropoiesis from adult peripheral blood, we observed modest stabilization of γ-globin mRNA by ZCCHC6 knock-out, suggesting that the ZCCHC6 machinery is active in adult erythropoiesis. Strategies to reactivate and stabilize γ-globin in adult red blood cells remains to be further optimized.

Akin to human β-globins, human α-globin mRNAs are selectively stabilized during terminal erythropoiesis, by a mechanism that entails binding of the αCP complex at polypyrimidine-rich sites within the 3’ UTR ^69^. In α^Constant^ ^Spring^ ^(CS)^ thalassemia, mutation of the 3’ UTR disrupts αCP complex formation, leading to accelerated mRNA decay, characterized by shortened poly(A) tails ^70^. Upon *Zcchc6* KO (but not in *Dis3l2* KO), RNA-seq results show that the embryonic α-like ζ-globin gene (*Hba-x*) was significantly upregulated in terminal erythroid cells at E18.5. It is therefore likely that ζ-globin and *γ*-globin share the RNA editing machinery that regulates their mRNA stability post-transcriptionally, most notably ZCCHC6, warranting future studies to investigate the consequences of ZCCHC6 KO in α^Constant^ ^Spring^ ^(CS)^ thalassemia.

## METHODS

### Animal Models

All experiments were performed with previously reported KO models for *Lin28a* and *Lin28b*, as well as KO and flox lines for *Zcchc6* and *Zcchc11*^13,71,72^. Whole body and conditional *Dis3l2* KO mice were generated in-house as follows: Targeted mESCs (Dis3l2^tm1a(EUCOMM)Hmgu^) were obtained from Helmholtz Zentrum München via the European Conditional Mouse Mutagenesis Program (EUCOMM)^73^. The gene-targeted ESCs were injected into chimeric blastocysts. Once injected, they were implanted into pseudopregnant Balb/c females and mutant mice were established. To ascertain germline transmission, we crossed chimeras to WT mice, screened the offspring, and obtained several Dis3l2 transgenic animals with germline transmission, after which the strain was expanded. Constitutive knockout of Dis3l2 was generated by crossing females carrying *Vasa-Cre* allele (Jackson laboratory, stock #006954) to *Dis3l2* transgenic males. The cross resulted in the constitutive deletion of Dis3l2 in all tissues by virtue of *Cre* expression in oocytes^74^. Conditional *Dis3l2* flox line was generated by crossing *Dis3l2* transgenic mice to Actin-Flippase (Flp) first (B6.Cg-Tg(ACTFLPe)9205Dym/J, the Jackson Laboratory Strain #005703) to remove Lac-Z reporter cassette, followed by *Cre* breeding.

The *Zcchc6* and *Dis3l2* flox animals were crossed with previously published *EpoR-Cre* mice to generate an erythroid-specific conditional KO mouse ^38^. Humanized beta-globin YAC mice were bred with *Zcchc6/Dis3l2* KO mice ^45^. Embryonic and adult ear tissue genotyping was performed either manually or automatically (using Transnetyx) to confirm the genotypes of KO, flox, and humanized lines. Conditional *Zcchc6* and *Dis3l2* KOs in the newly generated murine models were validated by western blots (1:1000 for both antibodies, ⍺-ZCCHC6; Protein Tech 25196-1-AP and ⍺-DIS3l2; NBP1-84740) on the sorted erythroid cells (PI-Hoechst-TER119+ cells). Animal experiments were performed by protocols approved by the Institutional Animal Care and Use Committee at Boston Children’s Hospital.

### Murine Blood/Tissue Collection and Analysis

Embryonic blood was collected from embryos harvested via Caesarian section. Blood of postnatal and adult mice was harvested through tail clipping or retro-orbital bleeding. During time-series analysis across various developmental ages, we bled each animal only once to avoid reticulocytosis secondary to anemia. Blood was collected using heparinized capillary tubes into EDTA-coated collection tubes, then subsequently processed using a HEMAVET machine and Fluorescence-Activated Single Cell Sorting (FACS) profiling.

One ul of blood was diluted into 50 ul of a buffered saline citrate glucose solution (BSCG) and mixed with equal parts of staining buffer (PBS + 2% serum + 1X pen/strep) containing a 1:100 dilution of each antibody: TER119 (BD Pharmingen 557909), CD71 (BD Clone: C2), Thiazole Orange (Sigma-Aldrich-390062), Hoechst 33342 (Invitrogen R37165), and Propidium Iodide (BD Pharmingen 51-66211E). All acquisition was performed on an LSR/FORTESSA cytometer, and the results were analyzed using FlowJo and Prism software. Spleen and liver cells were dissociated in ice-cold PBS, filtered, and then subjected to FACS analysis using the same staining panel.

For hemolytic marker assays, we collected plasma and serum samples, incubating the latter at room temperature for 30 minutes to allow for coagulation. We performed centrifugation at 1,500x g for 10 minutes at 4°C and collected plasma and serum samples for downstream analysis (western blots and blood biochemistry assays). The plasma was then mixed and heated in a protein preparation solution (RIPA buffer with protease and phosphatase inhibitors, and DTT). The protein samples were loaded onto Bis-Tris gels, and the transferred membranes were probed with primary antibodies (1:500 and 1:1000 for each antibody respectively: ⍺-Haptoglobin; Life Technologies # 16665-1-AP and ⍺-IgG; ab190475). LDH (Abcam no. ab102526) and bilirubin (Sigma Aldrich # MAK126) assays were performed according to the manufacturers’ instructions. The LDH samples were harvested from heparin-coated collection tubes instead of the EDTA-coated tubes.

### Blood Histology

Blood smears were prepared on frosted slides and stained with May-Grünwald and Giemsa solutions (Sigma-Aldrich; MG500 and GS500). The slides were placed in the May-Grünwald solution for two minutes, then were subsequently incubated in a diluted Giemsa solution (1:25 with milliQ H2O) for 12 minutes and washed with autoclaved Milli-Q H2O. Reticulocytes were identified using methylene blue staining protocol (Sigma-Aldrich; R4132). The stained slides were coated (VECTASHIELD) and examined under a microscope.

### Transcriptomic (RNA-seq) and Transcript Analysis (RT-qPCRs)

Enucleated erythroid cells (PI^-^Hoechst^-^TER119^+^ cells) were sorted directly into concentrated TRIzol (TRIzol™ LS Reagent). Total RNA was isolated using a column assay with DNase treatment (Direct-zol MicroPrep, ZYMO). The quantity and quality of RNA were evaluated using a nanodrop machine and RNA screen tape. We constructed an RNA-seq library with high-quality RNA (Both 260/280 and 260/230 over 1.7 with RNA integrity number >7) after ribosomal RNA depletion. For regular gene expression analysis, adapter-trimmed reads from the sequencer were mapped to the mouse genome, quantified, and analyzed using seq data analysis tools (Cutadapt, Bowtie, TopHat, HTSeq, R, and edgeR). For RT-qPCRs, cDNA was reverse transcribed from the isolated RNA (using the Maxima First Strand cDNA Synthesis Kit, Thermo Fisher Scientific). Then, quantitative PCR (qPCR) was performed using either SYBR Green or TaqMan assays (on a QuantStudio machine, Applied Biosystems).

### Human *In vitro* Erythropoiesis from Hematopoietic Stem/Progenitor Cells

Hematopoietic stem and progenitor (CD34+ HSPCs) cells derived from umbilical cord blood or adult peripheral blood were purchased (mixed donor vials, ALLCELLS/STEMCELL technologies) and subjected to the *in vitro* RBC differentiation protocol described in the previous study ^46^. Genetic KOs were generated in CD34+ HSPCs via nucleofection of CRISPR/Cas9 ribonucleoproteins (RNPs)^48^. First, CD34+ cells were thawed dropwise and cultured for two days before nucleofection in SFEM II media (StemCell Technologies, 09605) supplemented with 100 ng/mL SCF, TPO, FLT3L, and IL-6. The CD34+ HSPCs were nucleofected with Cas9 RNPs (125 pmol of sgRNA complexed with 105 pmol of Cas9 protein) using a Lonza 4D-Nucleofector. Cells were subsequently cultured for recovery and expansion, followed by the initiation of *in vitro* differentiation assays, and then harvested for genomic DNA extraction using a lysis buffer. The gRNA-targeted genomic loci (∼200 bp flanking regions of the expected Cas9 cut site) were PCR-amplified, gel-purified, and sanger-sequenced. The frameshifting Indel frequencies for each gRNA were quantified using TIDE analysis (http://tide.nki.nl). Hydroxyurea (Sigma) was purchased, freshly resuspended in ultra-pure water each time, and added to the cell cultures. For downstream qPCR analysis, cells were washed twice with ice-cold PBS, lysed in TRIzol, and their cDNAs were subsequently subjected to qPCRs. For Western blot and FACS analysis, we used the same method described in the murine sample analysis, except for the use of specific human antibodies: Hemoglobin β/γ/δ/ε Antibody (sc-390668) for western blot, and anti-Human CD71 (Fisher Scientific, BDB555537) and GlyA (CD235a, Beckman Coulter, A71564) for FACS analysis.

### Statistical Analysis

Data are expressed as mean ± SD. For hematological profile analysis, adjusted p-values were calculated from a linear mixed-effects model adjusted for sex (fixed effect) and litter information (random effect), followed by post-hoc multiple pairwise comparisons based on Tukey’s Honest Significant Difference (HSD) adjustment. For all other analyses, p-values were calculated using t-tests (* <0.05, ** <0.01, *** <0.001, **** <0.0001).

## Data availability

The resulting RNA-seq data is available on the Gene Expression Omnibus (GEO) database (GSE151184).

## Code availability

All scripts and codes are available upon request.

## Acknowledgements

The authors would like to thank Dr. Ursula Klingmüller for granting permission to use the *EpoR-cre* mouse model; Dr. Susanna Porcu for permission to use the Human *β-Globin YAC* mouse model; Dr. Dónal O’Carroll for sharing the *Zcchc6* and *Zcchc11* whole-body and conditional knockout mouse models; Dr. Vijay Sankaran for providing the globin primer lists and engaging in scientific discussions; current and former members of the Daley, Thorsten, and North labs for valuable discussions and technical assistance; the Harvard Medical School Biopolymers Facility for sequencing; and the Research Computing Group for providing the O2 High-Performance Compute Cluster. This work was supported by PCTC (Progenitor Cell Translational Consortium; NHLBI UO1-HL134812) and T32 grants from the National Heart, Lung, and Blood Institute (NHLBI), U54DK110805: A Center of Molecular Hematology from National Institute of Diabetes and Digestive and Kidney Diseases (NIDDK), as well as an early career grant from the National Blood Foundation.

## Author contributions

A.H. conceived the research question, performed experiments, and wrote the manuscript. A.Y. designed and conducted the Dis3l2 mice experiments. M.A.T.N. performed CRISPR/Cas9 knockouts and analyzed genome editing data. D.P. designed and established the ZCCHC6/11 mice study. Y.F. performed the blastocyst injections for the Dis3l2 study and guided the EpoR-cre and YAC mice studies. M.B. conducted mouse genotyping and RNA experiments. C.K. designed and provided guidance on FACS profiling. T.N. advised on the study. V.L. set up the human *in vitro* erythropoiesis study and revised the manuscript. S.O. designed the EpoR-cre and YAC mice studies and advised on the study. G.Q.D. conceived the idea, supervised the study, and wrote the manuscript.

---

**Extended Data Fig. 1.**
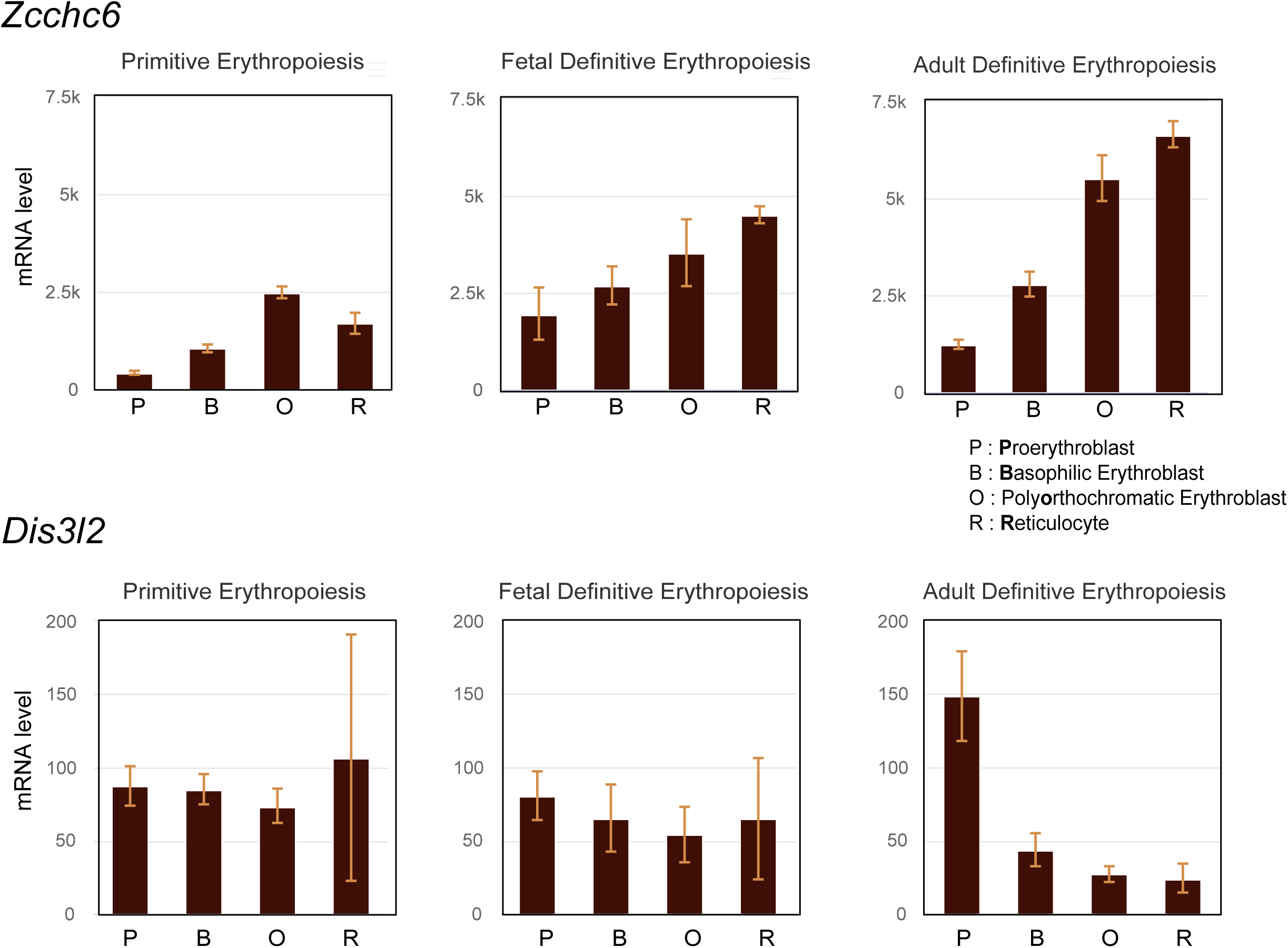
Expression Profiles of Zcchc6 and Dis3l2 mRNAs During Erythropoiesis. (from the Erythron Database). Expression Levels of Zcchc6 and Dis3l2 mRNAs During Mouse Primitive and Definitive (fetal liver and adult bone marrow) Erythropoiesis.

**Extended Data Fig. 2.**
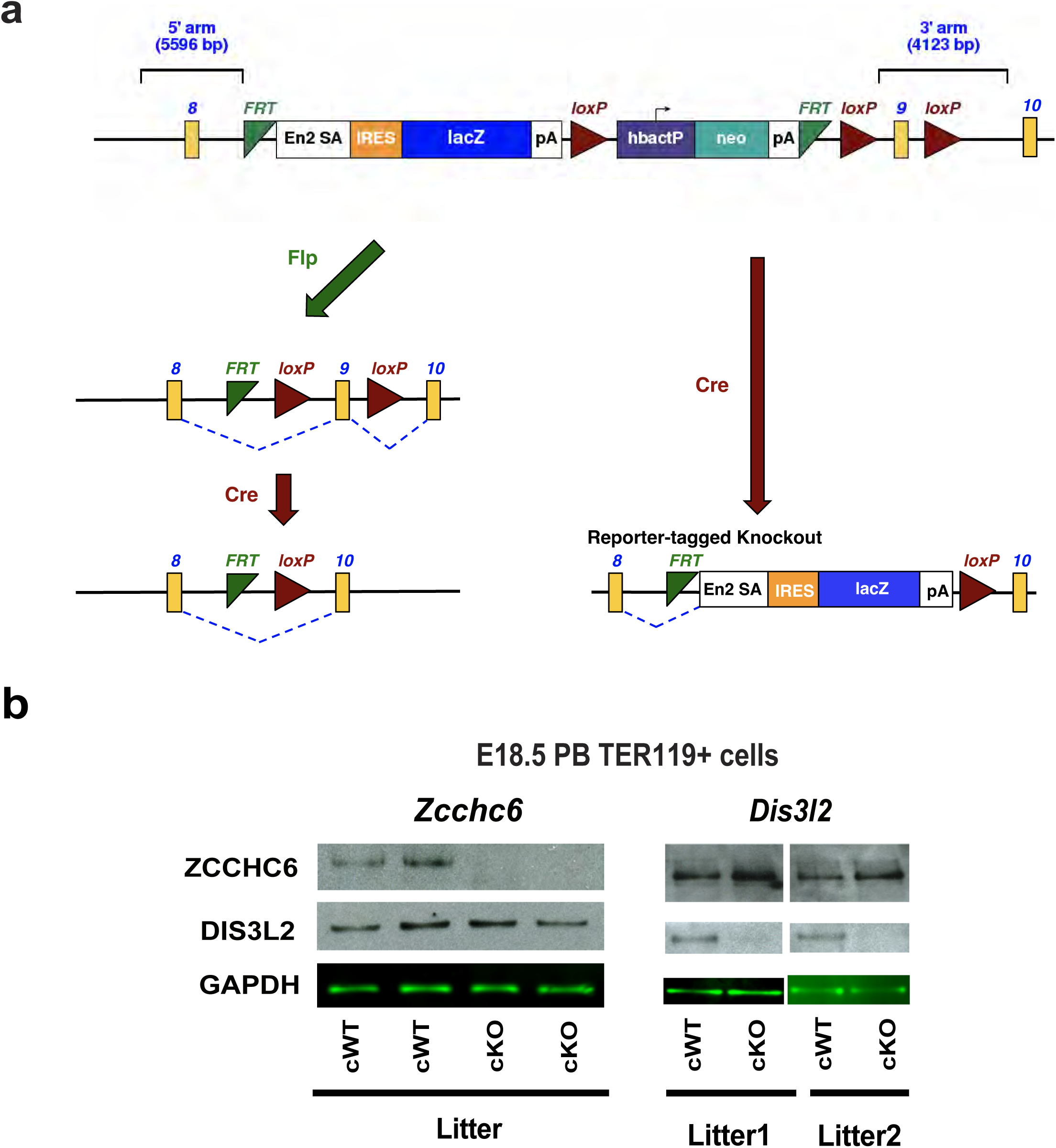
Allele Map of the Dis3l2 Knockout Model and Conditional Knockout Validation. **(a)** Allele Map of the *Dis3l2* Knockout Models. **(b)** Western Blots of Erythroid Cells from Embryos Carrying only *floxed* (cWT) and both *floxed* and *EpoR-Cre* (cKO) Alleles for Validation.

**Extended Data Fig. 3.**
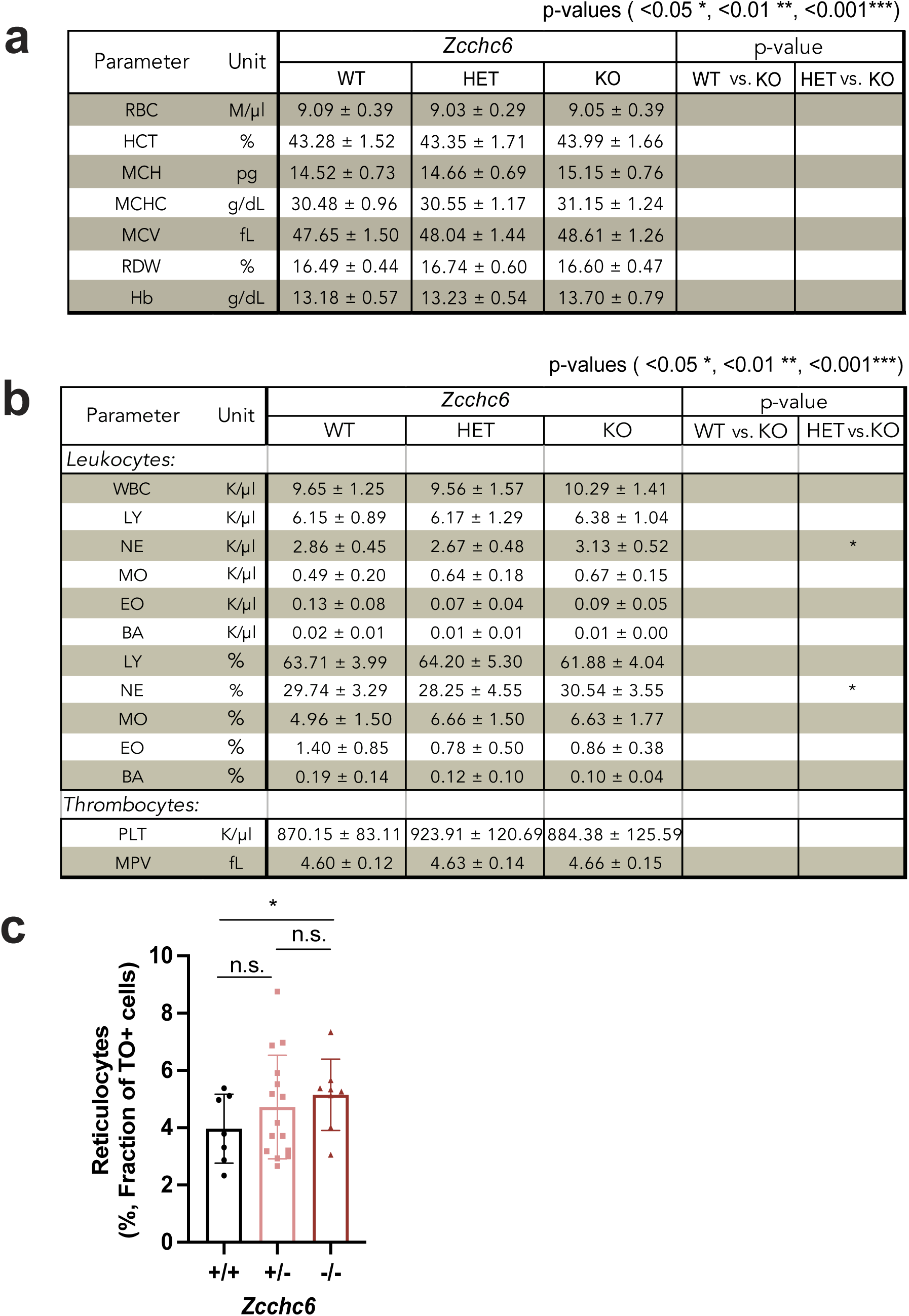
Analysis of Whole-Body *Zcchc6* Knockout Young Adult Mice (6–7 Weeks). **(a)** Hematological Profiles of Peripheral Blood from Retro-Orbital Bleeding (mean ± SD). HCT; Hematocrit, MCH; Mean corpuscular hemoglobin, MCHC; Mean corpuscular hemoglobin concentration, MCV; Mean corpuscular volume, RDW; Red cell Distribution Width. Multiple litters were used. **(b)** Leukocytes and Thrombocyte panels. LY; lymphocytes, NE; neutrophils, MO; monocytes, EO; eosinophils, BA; basophils, PLT; Platelet or thrombocyte count, MPV; Mean Platelet Volume. **(c)** Quantification of reticulocyte (TO+) population per genotype (mean ± SD).

**Extended Data Fig. 4.**
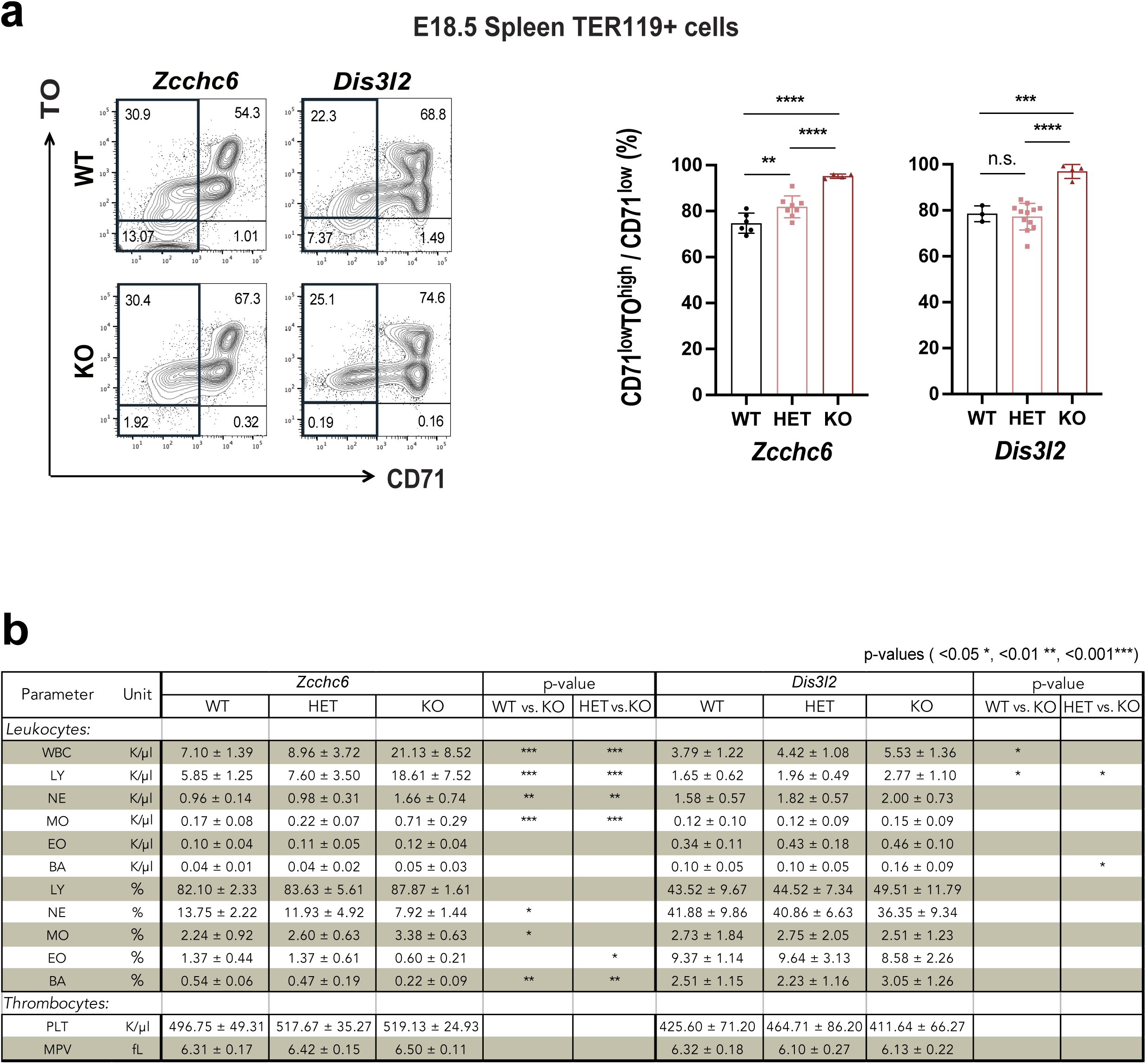
Additional Analysis of Whole-Body *Zcchc6/Dis3l2* Knockout Mice. (a, left figures) Flow cytometry profiles of erythroid cells from spleens. Representative profiles of each KO mouse and WT littermates were displayed. **(a, right graphs)** Quantification of reticulocyte population per genotype. **(b)** Hematological profiles in the PB of whole-body KOs (Leukocytes and Thrombocyte panels).

**Extended Data Fig. 5.**
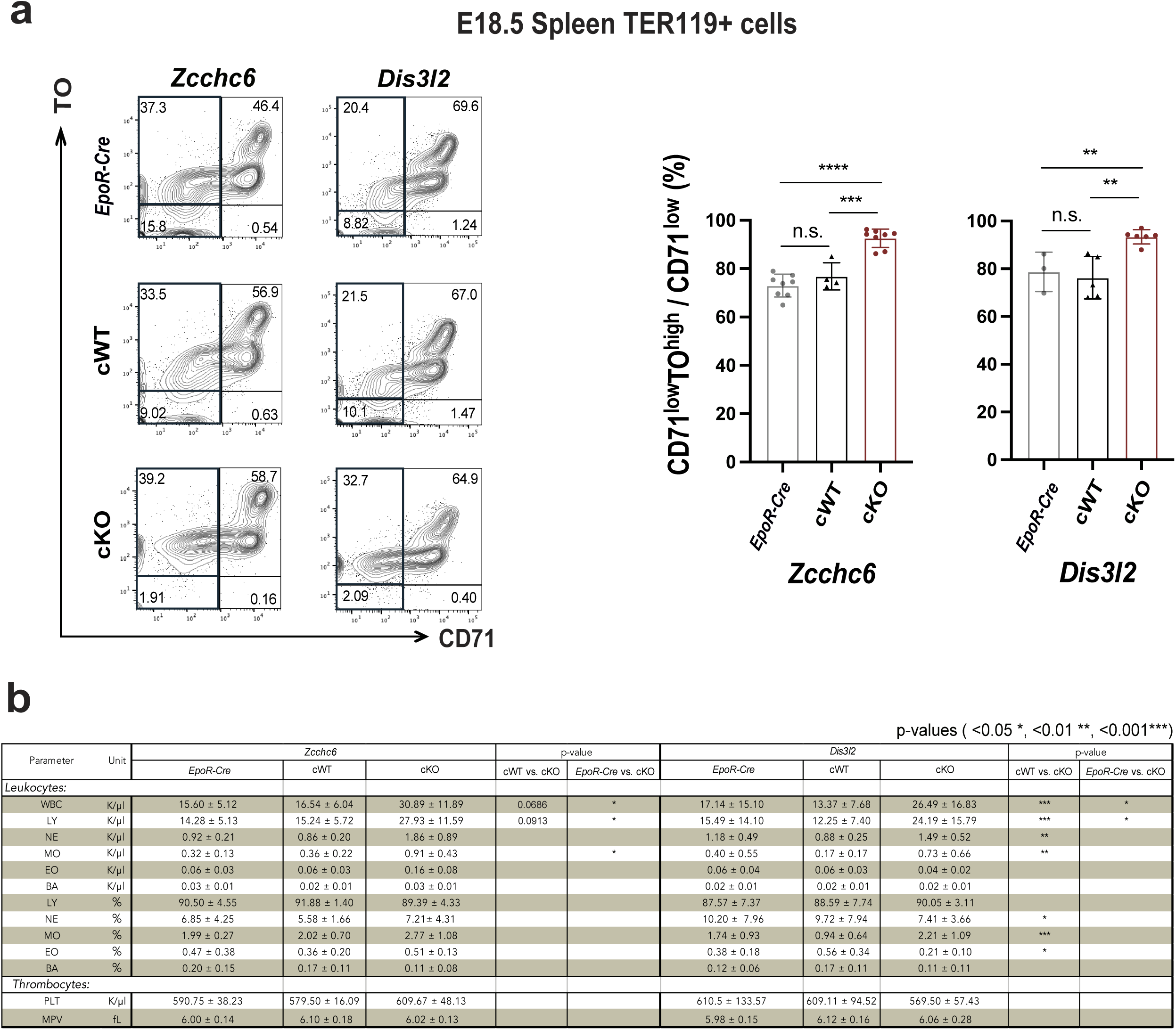
Additional Analysis of Conditional *Zcchc6/Dis3l2* KO. **(a)** Flow cytometry profiles of erythroid cells from the spleen. Representative profiles and reticulocyte quantification of embryos carrying only the *EpoR-Cre* allele, only the *floxed* allele; cWTs, and both alleles; cKOs. **(b)** Hematological profiles in the PB of conditional KOs. (Leukocytes and Thrombocyte panels).

**Extended Data Fig. 6.**
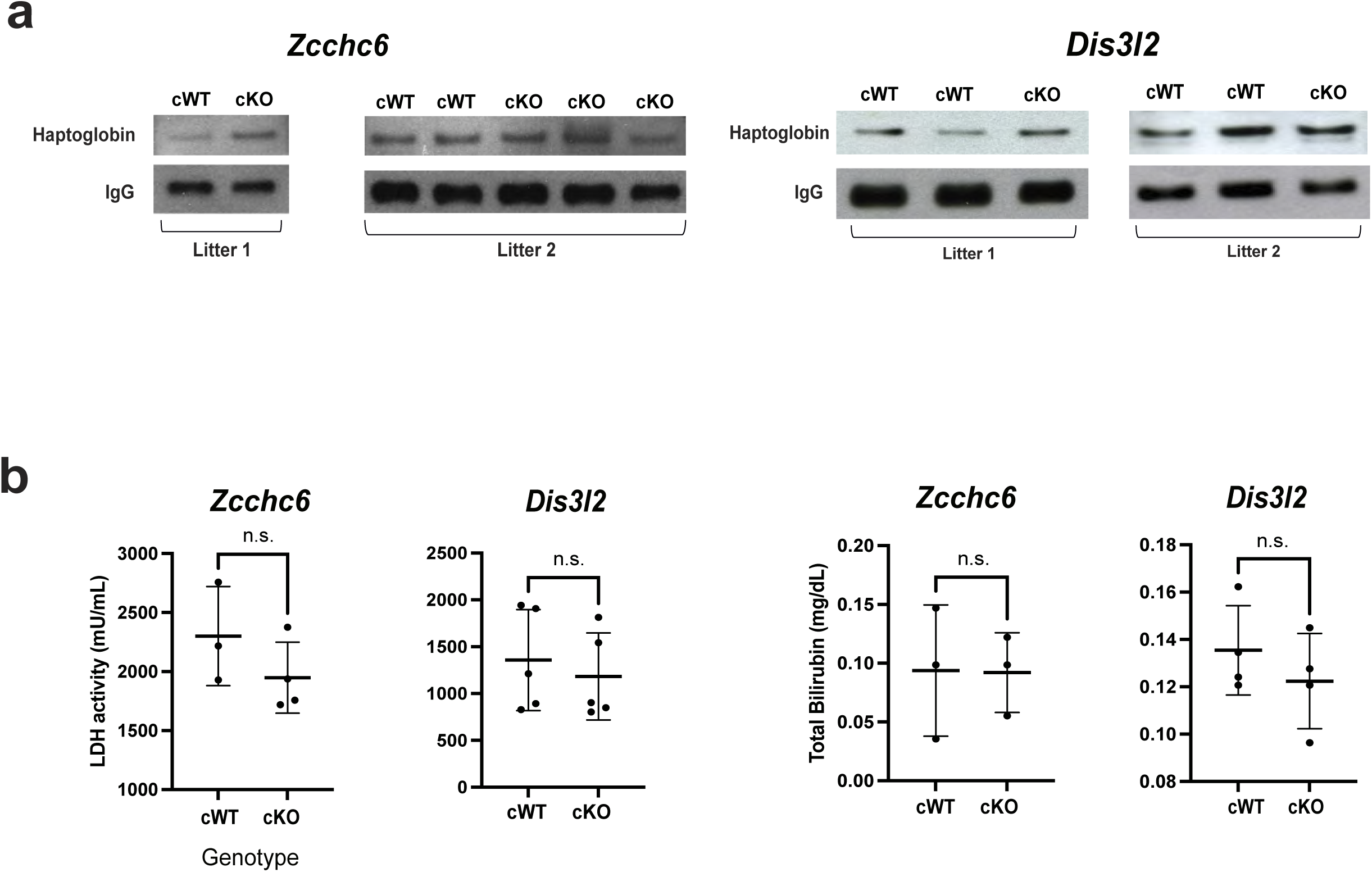
Analysis of Hemolytic Anemia Markers. **(a)** Western blots of Haptoglobin in cKOs and cWTs. **(b)** LDH and Bilirubin assay results from serum and plasma of cKO mice.

**Extended Data Fig. 7.**
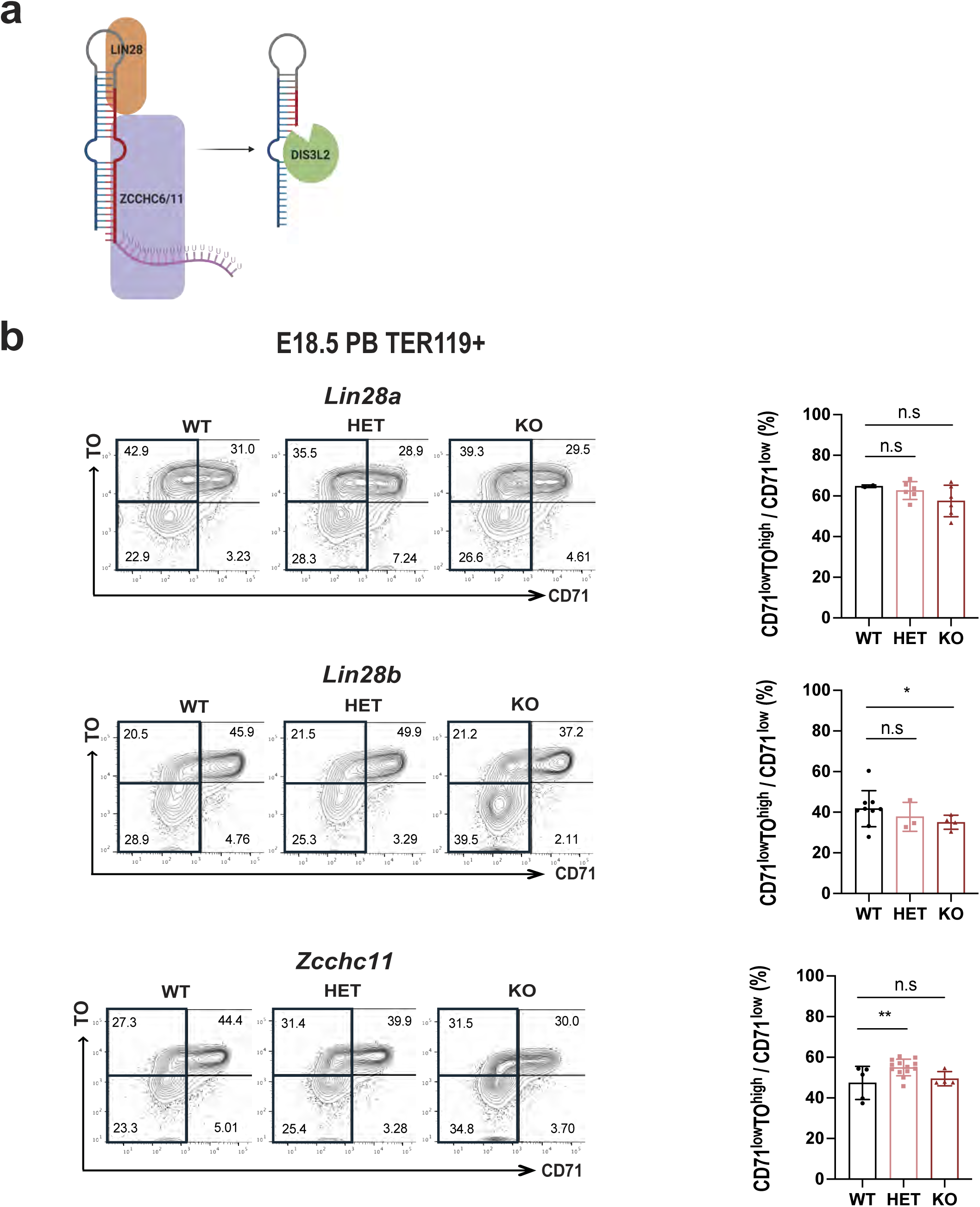
Characterizing the Impact of Knockouts in Pathway-Related Genes. **(a)** Previously published model for the RNA decay machinery (LIN28-ZCCHC6/11-DIS3L2) on the microRNA regulation. **(b)** FACS profiles and population quantification for each whole-body KO model (*Lin28a*, *Lin28b*, and *Zcchc11* sequentially).

**Extended Data Fig. 8.**
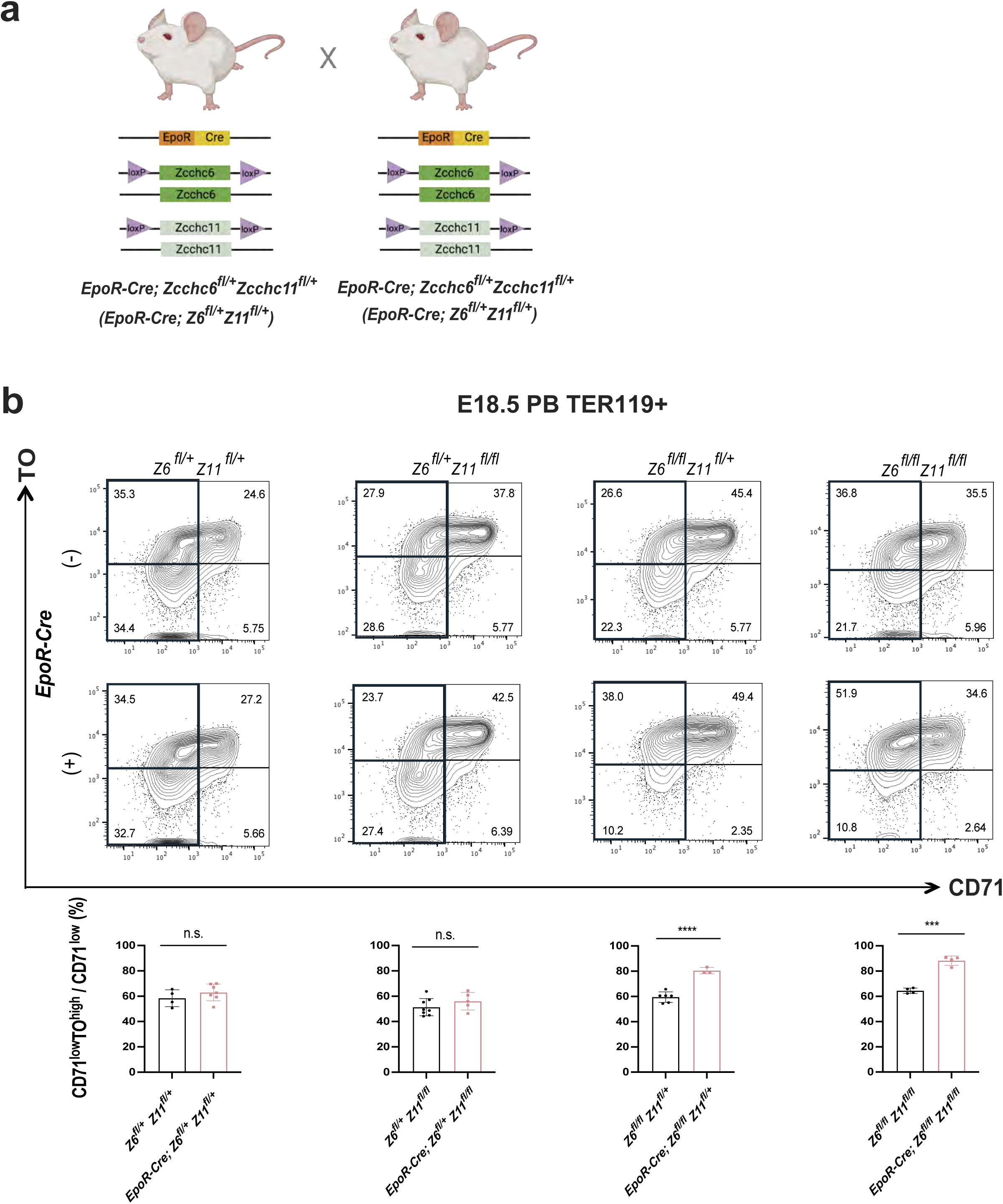
Analysis of *Quad-floxed* Conditional Knockouts. **(a)** *Quad-floxed* conditional KO breeding scheme for *Zcchc6* and its paralog *Zcchc11*. **(b)** Representative FACS profiles of the *quad-floxed* embryos and quantification of reticulocytes.

**Extended Data Fig. 9.**
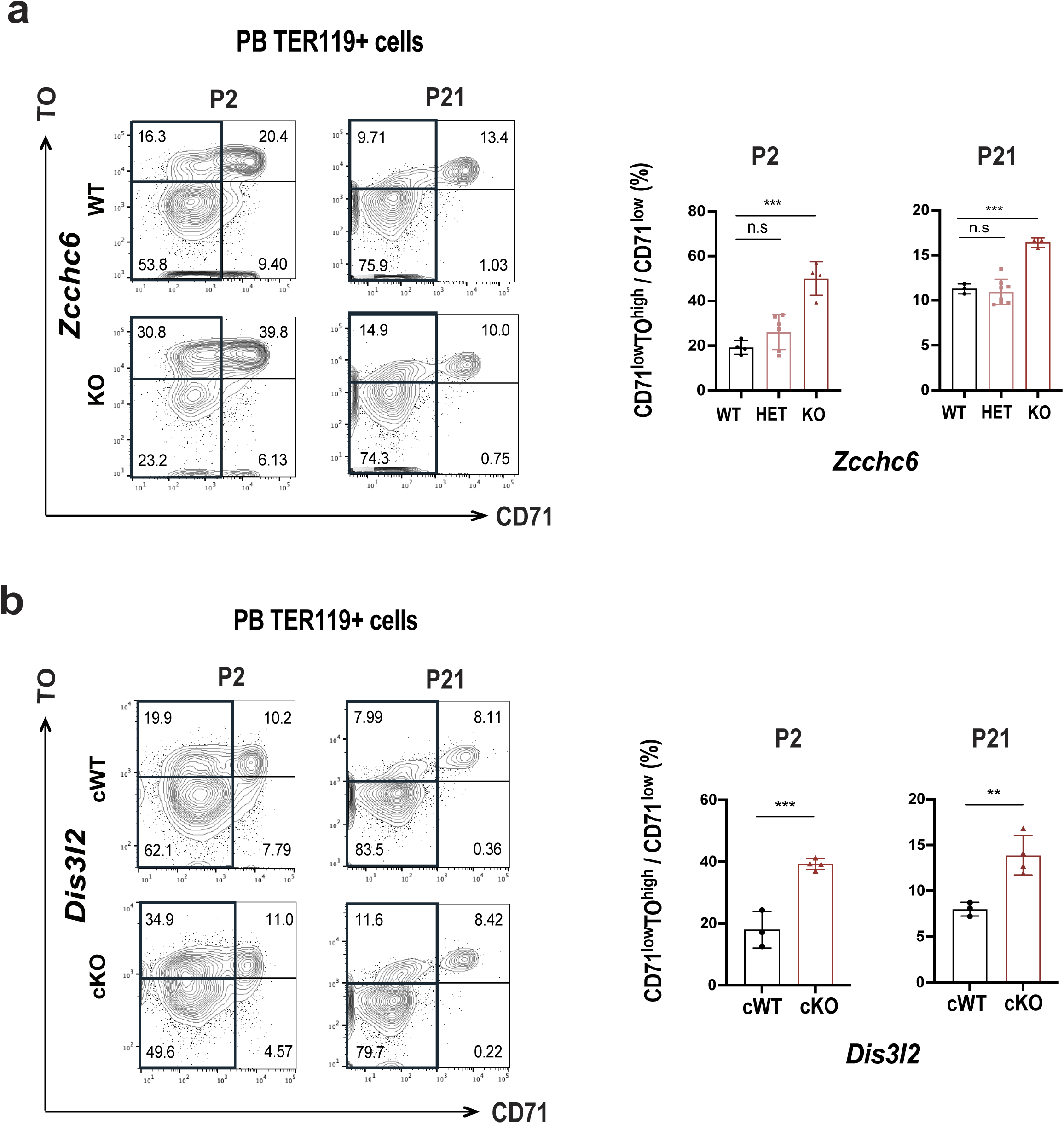
Postnatal Profiling of *Zcchc6* KO and *Dis3l2* cKO. **(a)** Postnatal PB FACS profiling and quantification of whole body *Zcchc6* KO model **(b)** and *Dis3l2* cKO model.

**Extended Data Fig. 10.**
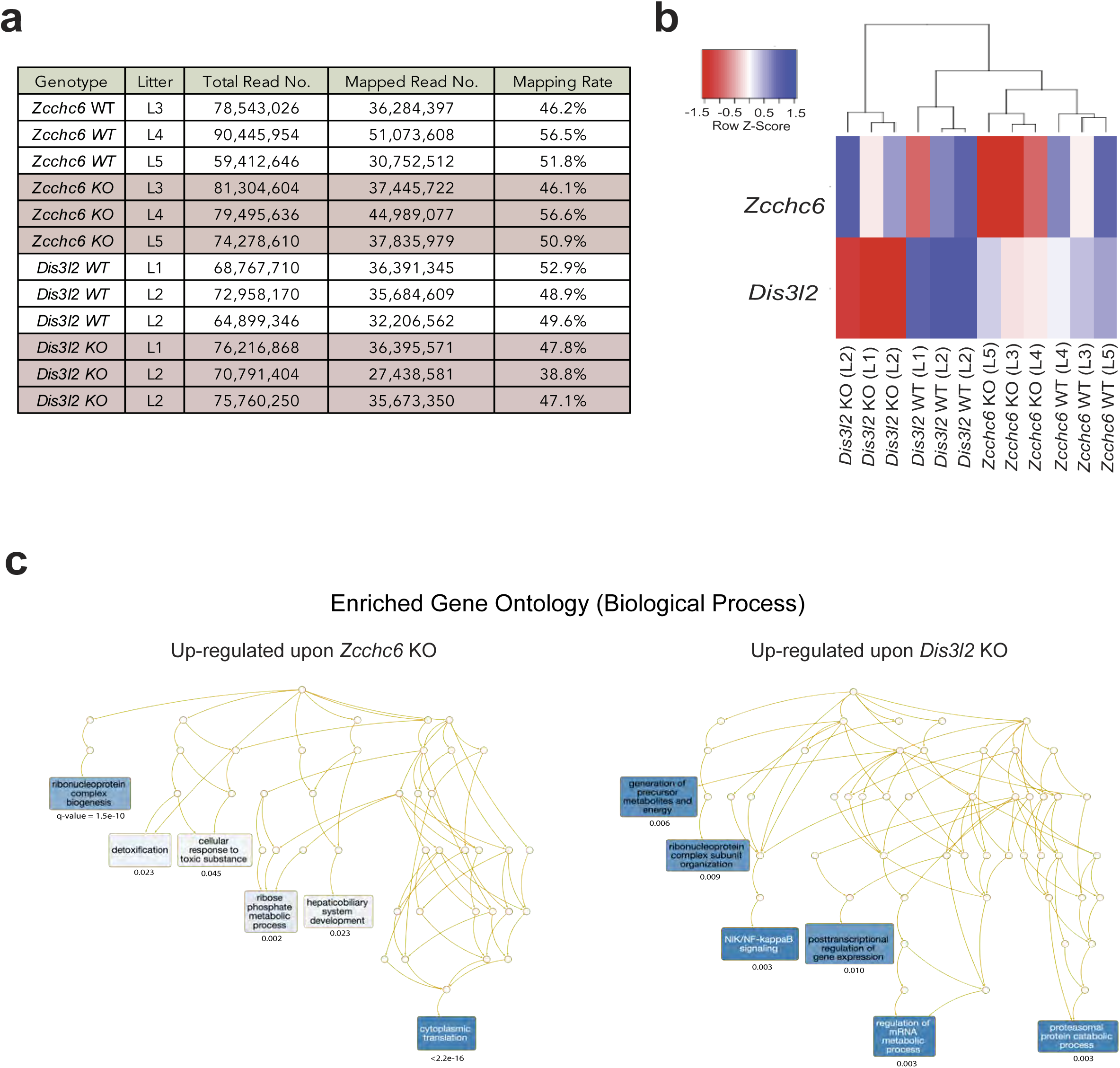
RNA-seq Data Analysis. **(a)** Depth and mapping rates of bulk RNA-seq experiments. **(b)** Hierarchical clustering of samples based on the expression of *Zcchc6* and *Dis3l2*. **(c)** Enriched gene ontology terms and their statistical significances for each KO model.

**Extended Data Fig. 11.**
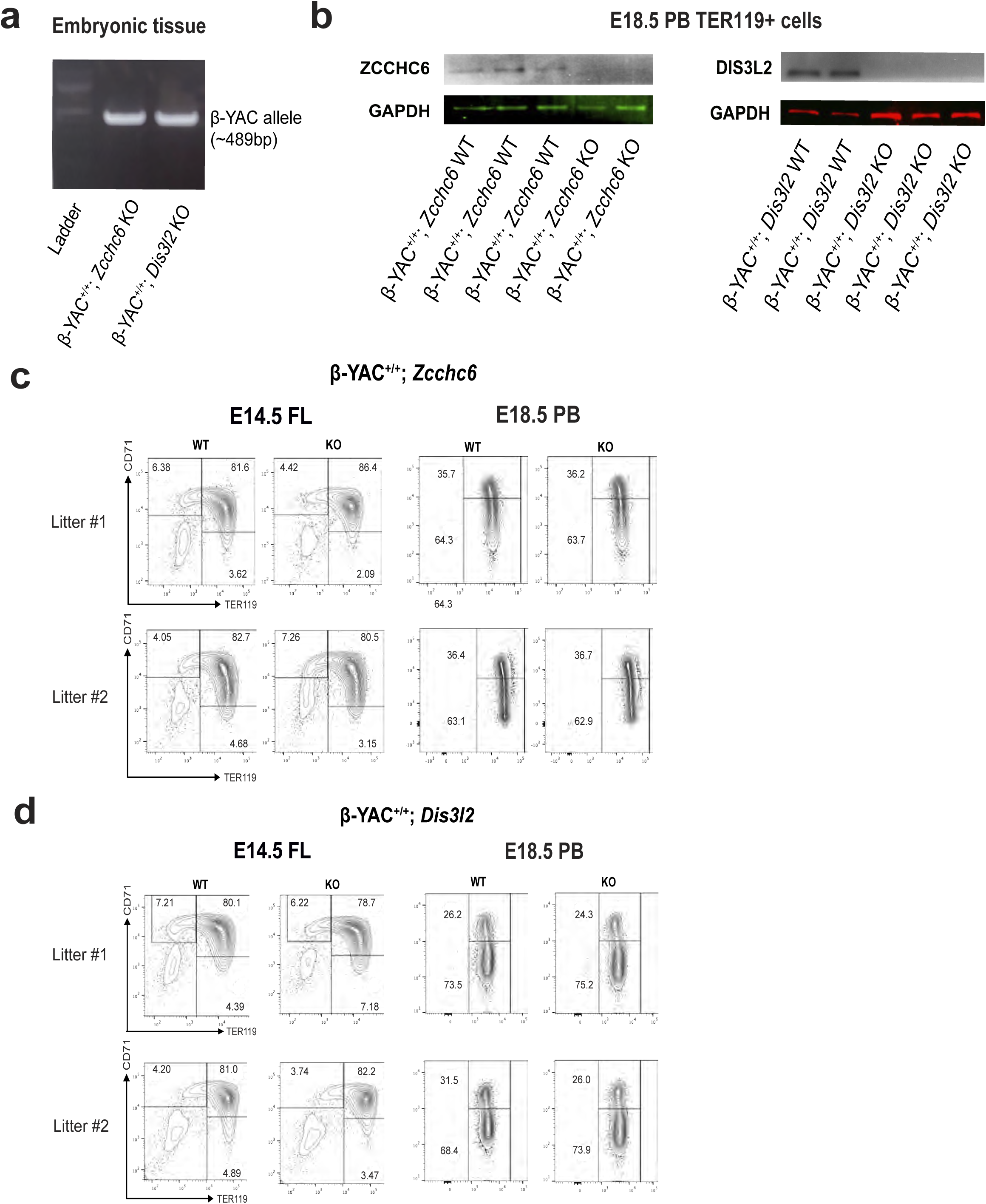
Analysis of Humanized Mice Upon *Zcchc6/Dis3l2* KO. **(a)** Genomic PCR analysis of the human β-YAC allele in offspring. **(b)** Validation of preserved *Zcchc6/Dis3l2* KO in the crossed humanized mice using western blots. **(c & d)** FACS profiles of erythroid maturation from E14.5 fetal liver and E18.5 PB of subject mice.

**Extended Data Fig. 12.**
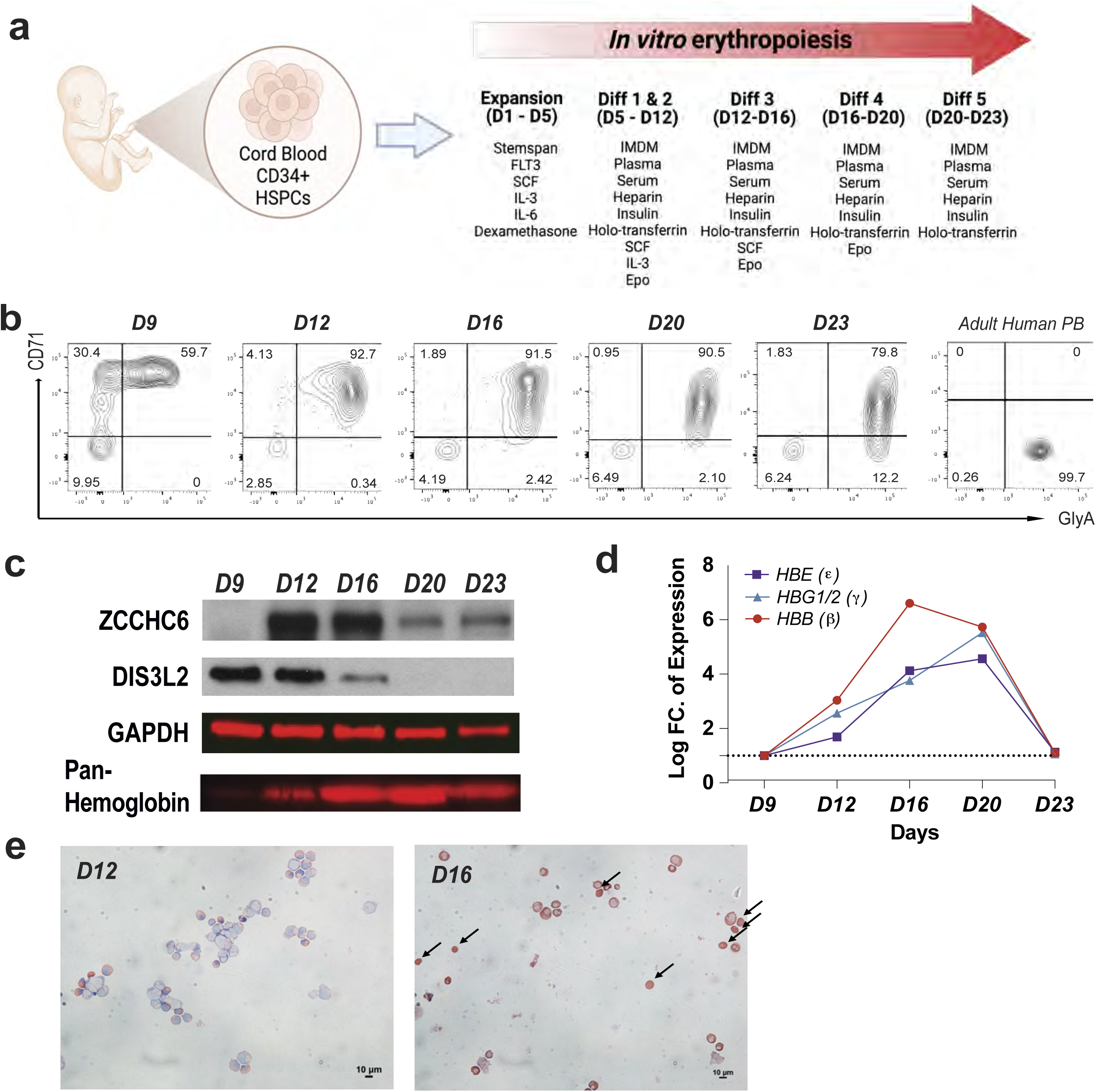
Human *In Vitro* Erythropoiesis. **(a)** Scheme of *in vitro* erythropoiesis from human cord blood-derived hematopoietic stem/progenitor cells (HSPCs; CD34+ cells). **(b)** Time-series FACS profiles to follow erythroid maturation status (Adult human PB served as a positive control). **(c)** Time-series protein expression profiles of ZCCHC6, DIS3L2, and hemoglobin during the *in vitro* erythropoiesis by western blots. **(d)** RNA expression profiles of β-like globins using qPCRs during the in vitro erythropoiesis. **(e)** Histology of cytospined cells (stained by May-Grünwald Giemsa method).

**Extended Data Fig. 13.**
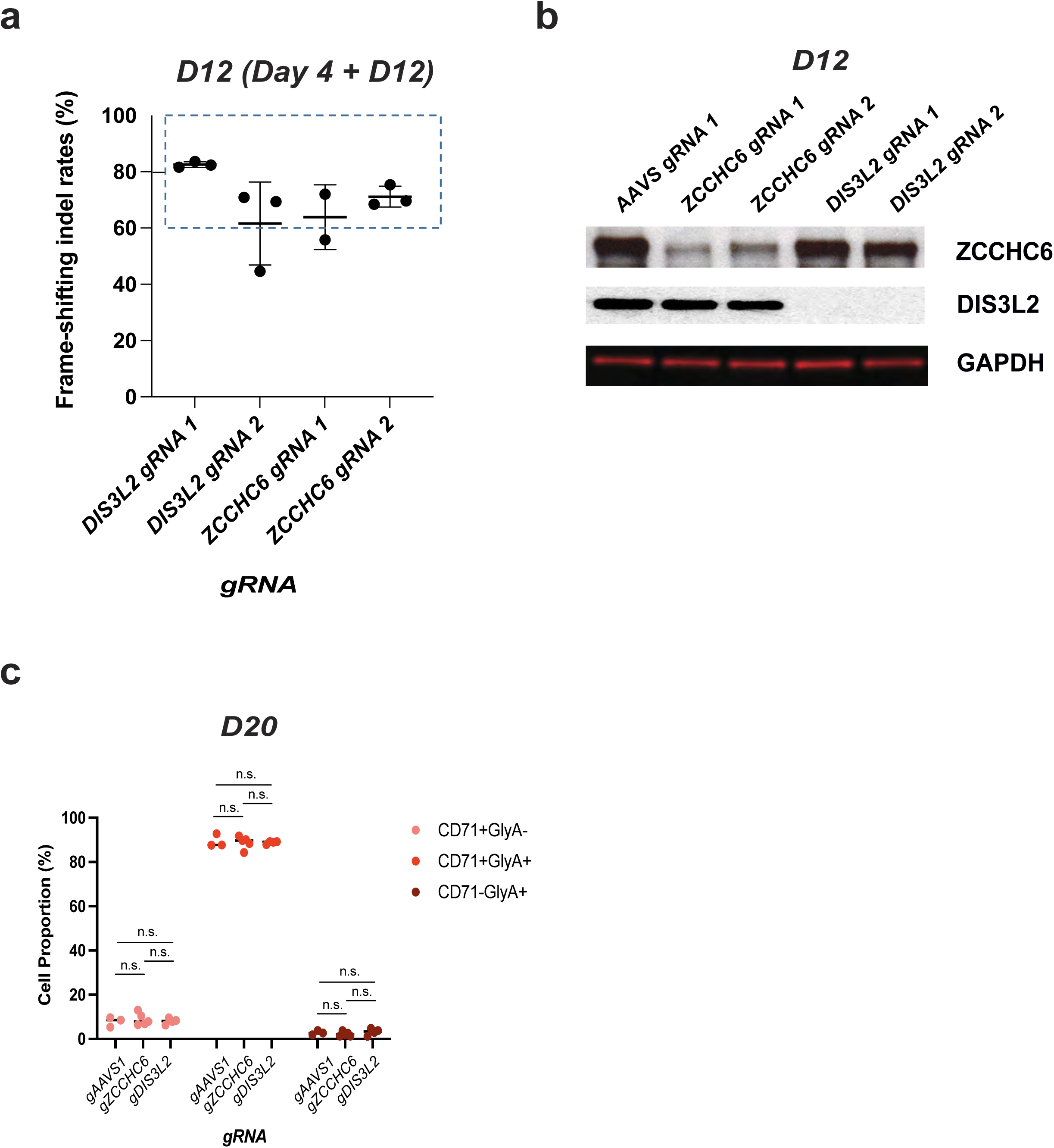
Analysis of CRISPR KO in Differentiated Erythroid Cells from CB. **(a)** Genomic editing and **(b)** Protein profiles to access efficiency of CRISPR/Cas9-mediated KOs. **(c)** Summary graph of erythroid maturation status by FACS (CD71 and GlyA) in CRISPR/Cas9 KOs.

**Extended Data Fig. 14.**
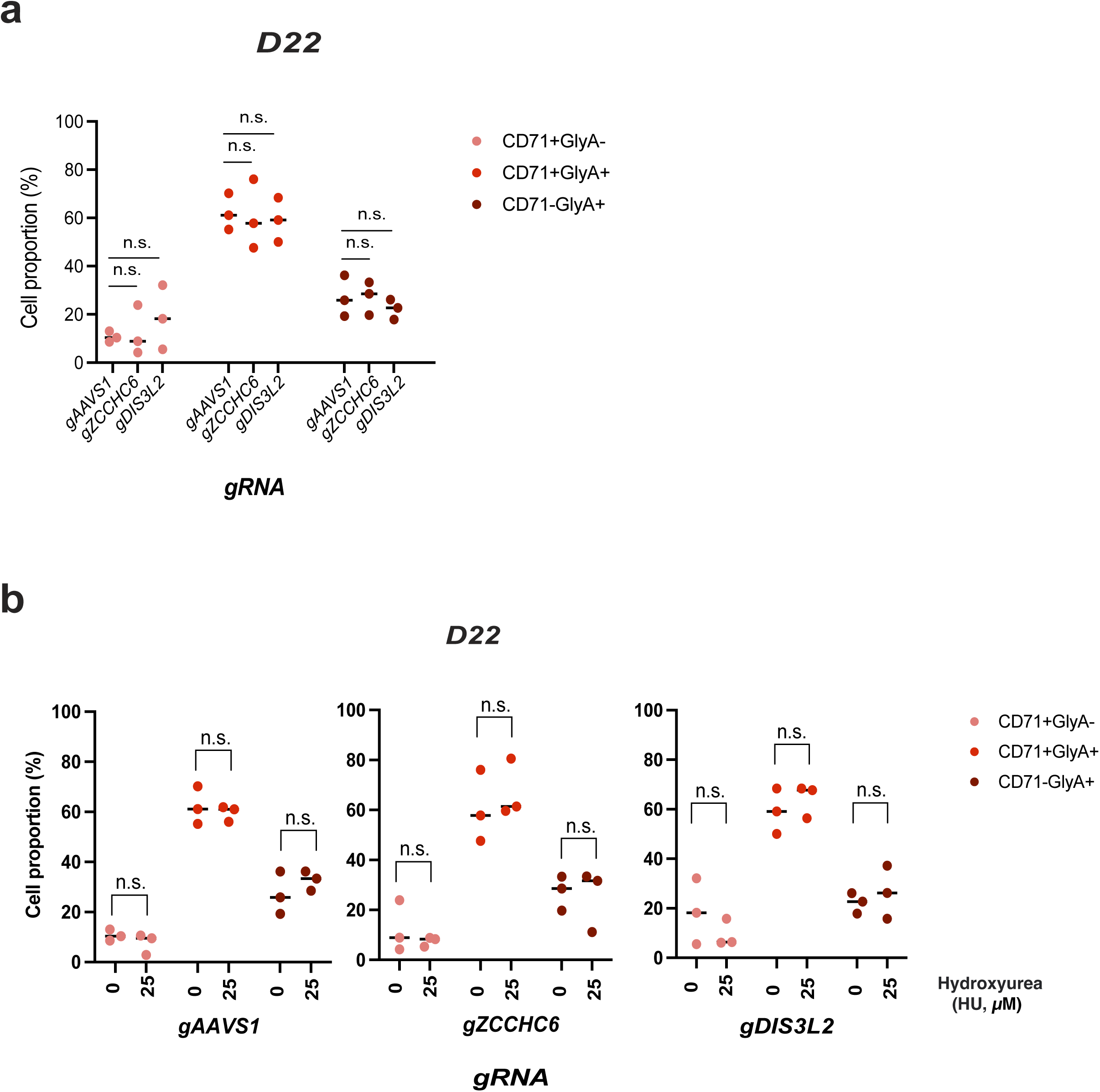
Analysis of CRISPR KO in Differentiated Erythroid Cells from PB Following HU treatment. Summary graph of erythroid maturation status by FACS (CD71 and GlyA) in **(a)** CRISPR/Cas9 KOs **(b)** CRISPR/Cas9 KOs & HU treatment

## SUPPLEMENTAL TABLES

**Supplemental Table 1.**
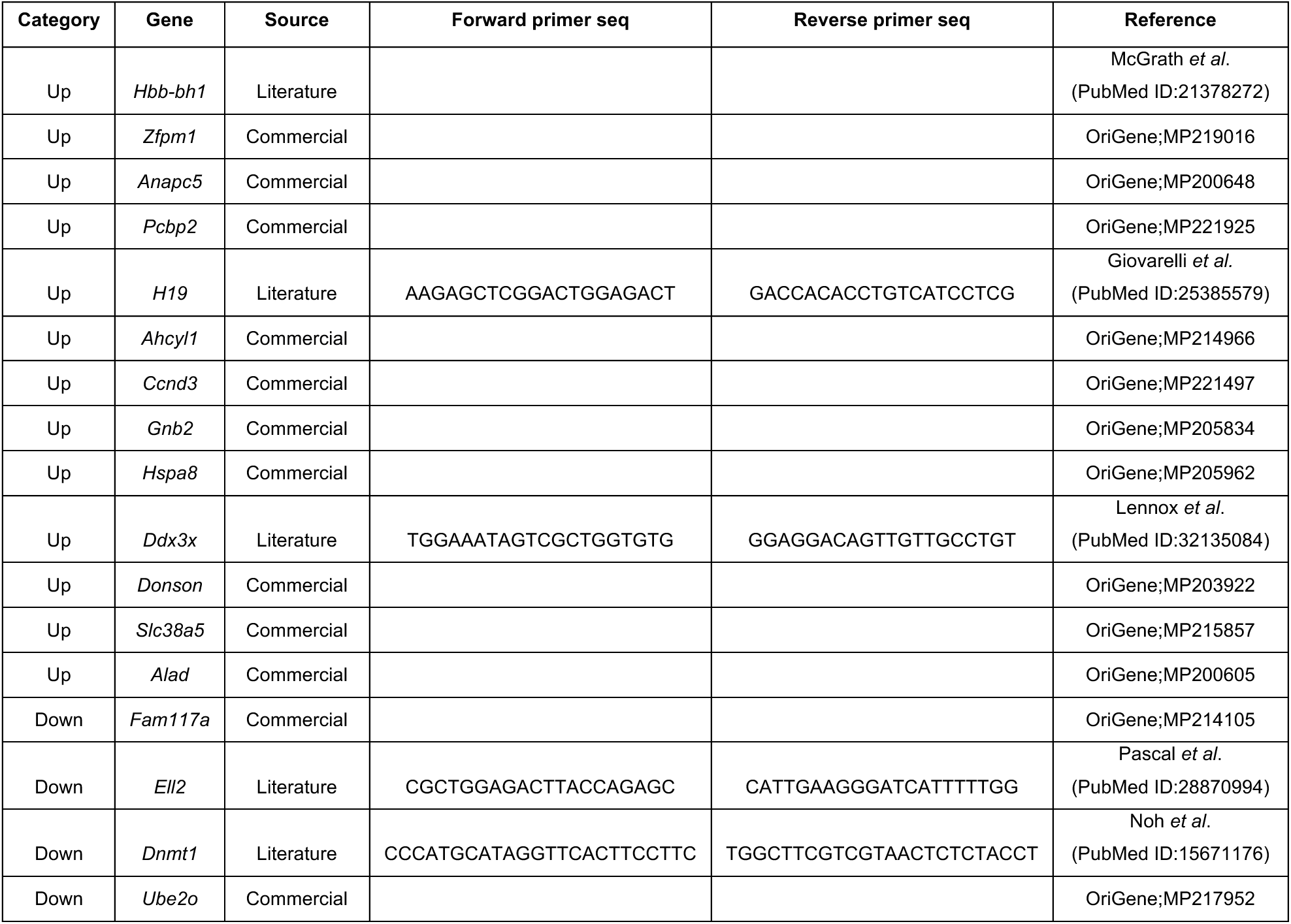
qPCR Primers for RNA-seq Validation.

**Supplemental Table 2.**
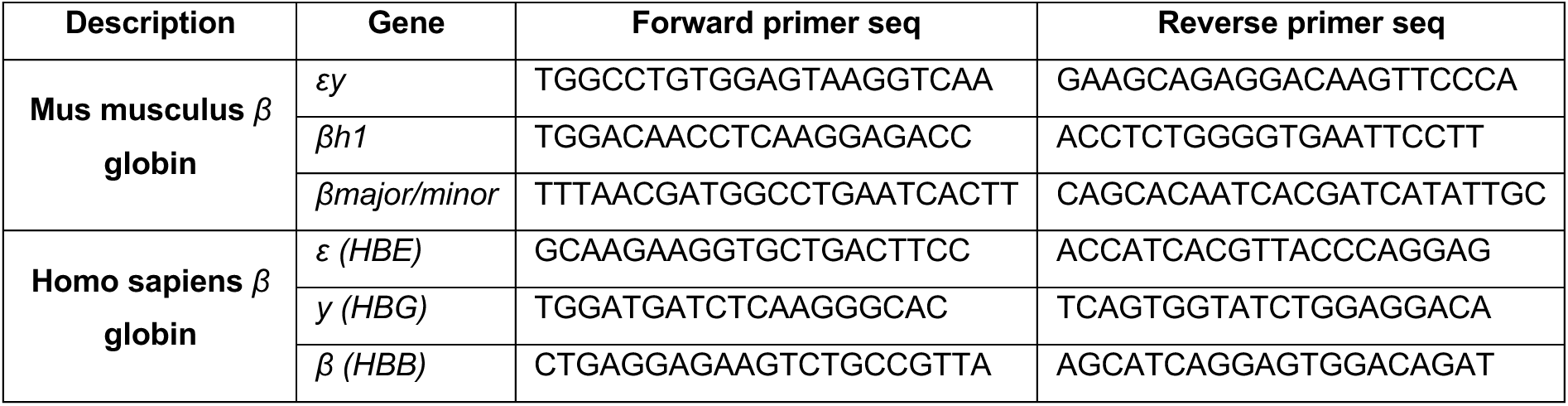
qPCR Primers for β-like globin profiling.

**Supplemental Table 3.**
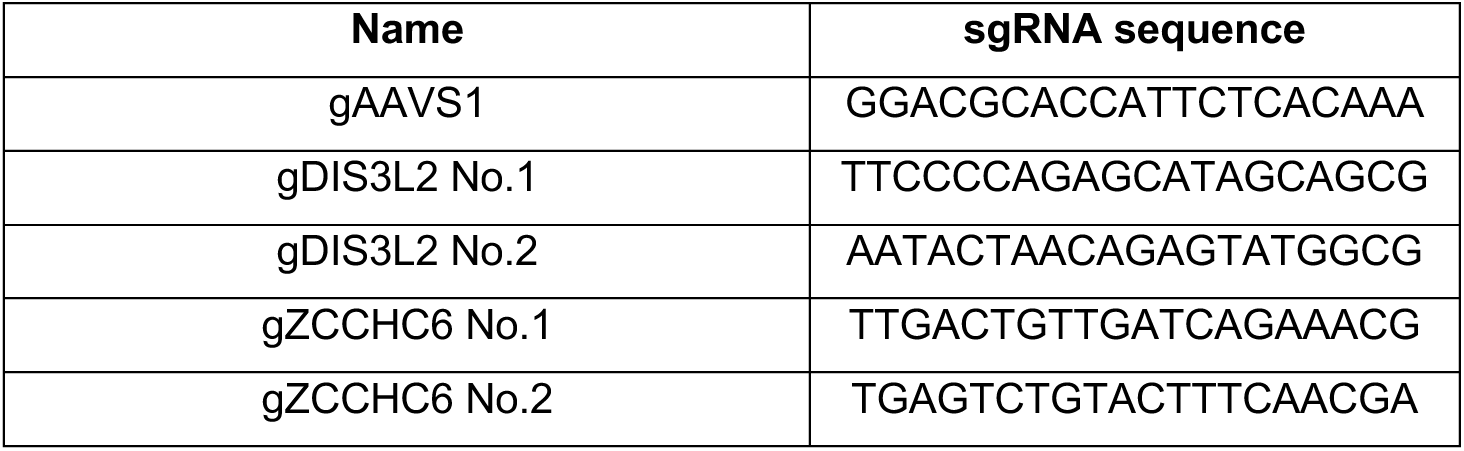
Guide RNA (gRNA) Sequences for CRISPR/Cas9 Knockout (KO)

