## Extended Data for "RNA Editors Sculpt the Transcriptome During Terminal Erythropoiesis" (from another version of this preprint: version 1, 2025-04-05)

### Extended Data Fig. 1

#### Expression Profiles of *Zcchc6* and *Dis3l2* mRNAs During Erythropoiesis.

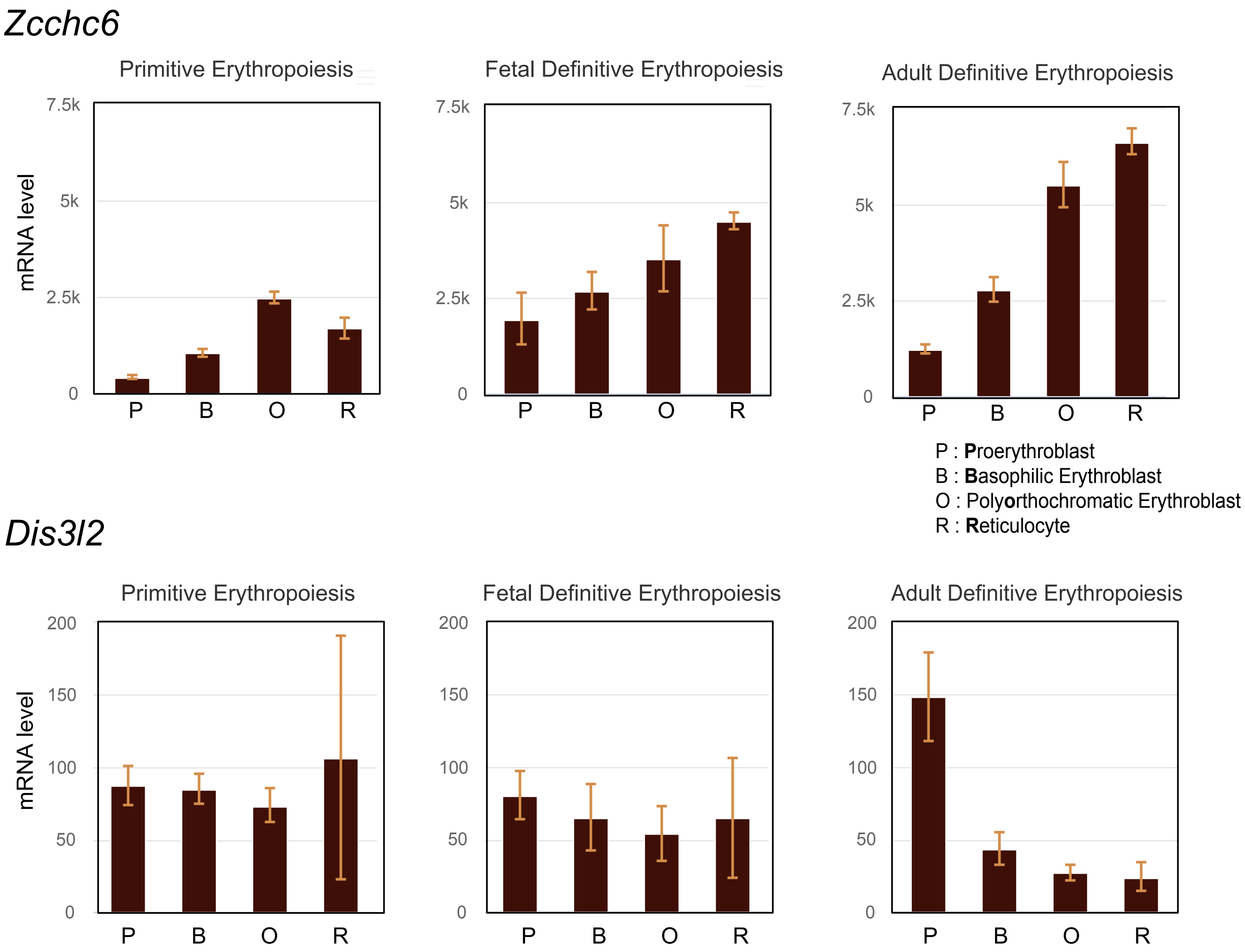

### Extended Data Fig. 2

#### Allele Map of the *Dis3l2* Knockout Model and Conditional Knockout Validation.

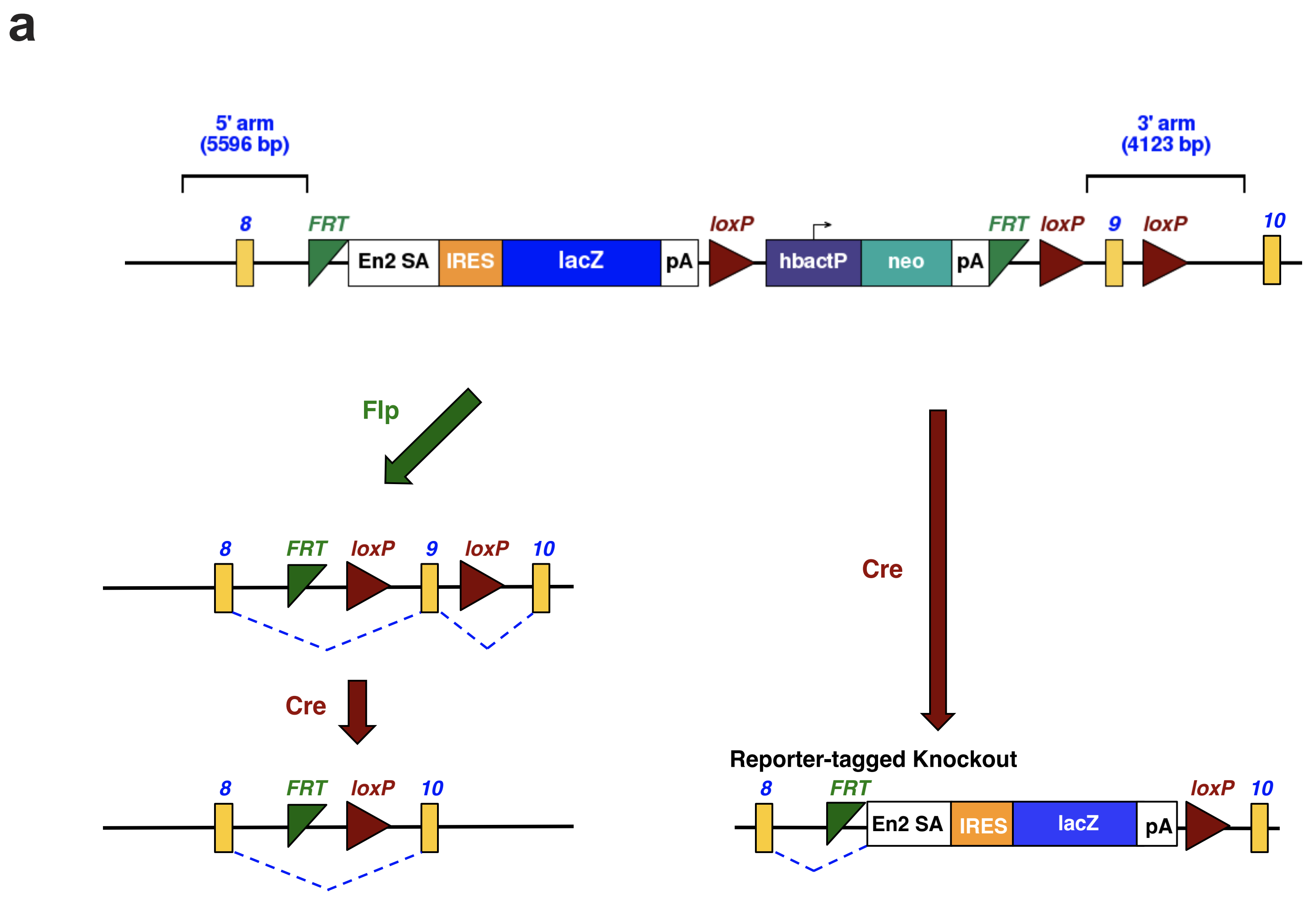


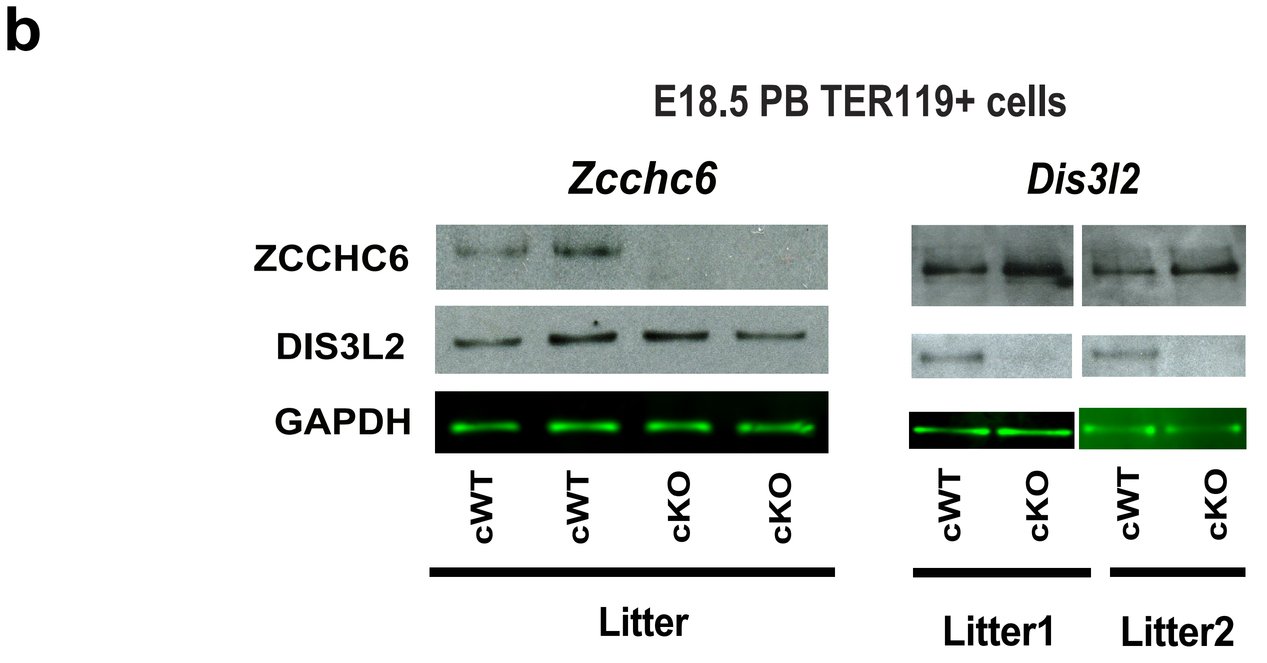

### Extended Data Fig. 3

#### Analysis of Whole-Body *Zcchc6* Knockout Young Adult Mice (6–7 Weeks).

**a**

p-values ( <0.05 *, <0.01 **, <0.001***)

| Parameter | Unit | *Zcchc6* | | | p-value | |
|---|---|---|---|---|---|---|
| | | WT | HET | KO | WT vs. KO | HET vs. KO |
| RBC | M/µl | 9.09 ± 0.39 | 9.03 ± 0.29 | 9.05 ± 0.39 | | |
| HCT | % | 43.28 ± 1.52 | 43.35 ± 1.71 | 43.99 ± 1.66 | | |
| MCH | pg | 14.52 ± 0.73 | 14.66 ± 0.69 | 15.15 ± 0.76 | | |
| MCHC | g/dL | 30.48 ± 0.96 | 30.55 ± 1.17 | 31.15 ± 1.24 | | |
| MCV | fL | 47.65 ± 1.50 | 48.04 ± 1.44 | 48.61 ± 1.26 | | |
| RDW | % | 16.49 ± 0.44 | 16.74 ± 0.60 | 16.60 ± 0.47 | | |
| Hb | g/dL | 13.18 ± 0.57 | 13.23 ± 0.54 | 13.70 ± 0.79 | | |

**b**

p-values ( <0.05 *, <0.01 **, <0.001***)

| Parameter | Unit | *Zcchc6* | | | p-value | |
|---|---|---|---|---|---|---|
| | | WT | HET | KO | WT vs. KO | HET vs.KO |
| *Leukocytes:* | | | | | | |
| WBC | K/µl | 9.65 ± 1.25 | 9.56 ± 1.57 | 10.29 ± 1.41 | | |
| LY | K/µl | 6.15 ± 0.89 | 6.17 ± 1.29 | 6.38 ± 1.04 | | |
| NE | K/µl | 2.86 ± 0.45 | 2.67 ± 0.48 | 3.13 ± 0.52 | | * |
| MO | K/µl | 0.49 ± 0.20 | 0.64 ± 0.18 | 0.67 ± 0.15 | | |
| EO | K/µl | 0.13 ± 0.08 | 0.07 ± 0.04 | 0.09 ± 0.05 | | |
| BA | K/µl | 0.02 ± 0.01 | 0.01 ± 0.01 | 0.01 ± 0.00 | | |
| LY | % | 63.71 ± 3.99 | 64.20 ± 5.30 | 61.88 ± 4.04 | | |
| NE | % | 29.74 ± 3.29 | 28.25 ± 4.55 | 30.54 ± 3.55 | | * |
| MO | % | 4.96 ± 1.50 | 6.66 ± 1.50 | 6.63 ± 1.77 | | |
| EO | % | 1.40 ± 0.85 | 0.78 ± 0.50 | 0.86 ± 0.38 | | |
| BA | % | 0.19 ± 0.14 | 0.12 ± 0.10 | 0.10 ± 0.04 | | |
| *Thrombocytes:* | | | | | | |
| PLT | K/µl | 870.15 ± 83.11 | 923.91 ± 120.69 | 884.38 ± 125.59 | | |
| MPV | fL | 4.60 ± 0.12 | 4.63 ± 0.14 | 4.66 ± 0.15 | | |

**c**

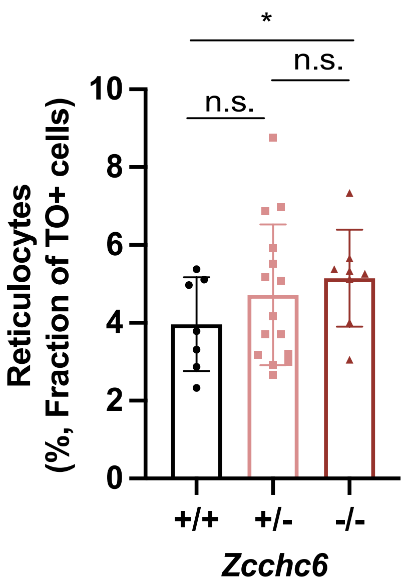

### Extended Data Fig. 4

#### Additional Analysis of Whole-Body *Zcchc6/Dis3l2* Knockout Mice.

**a**

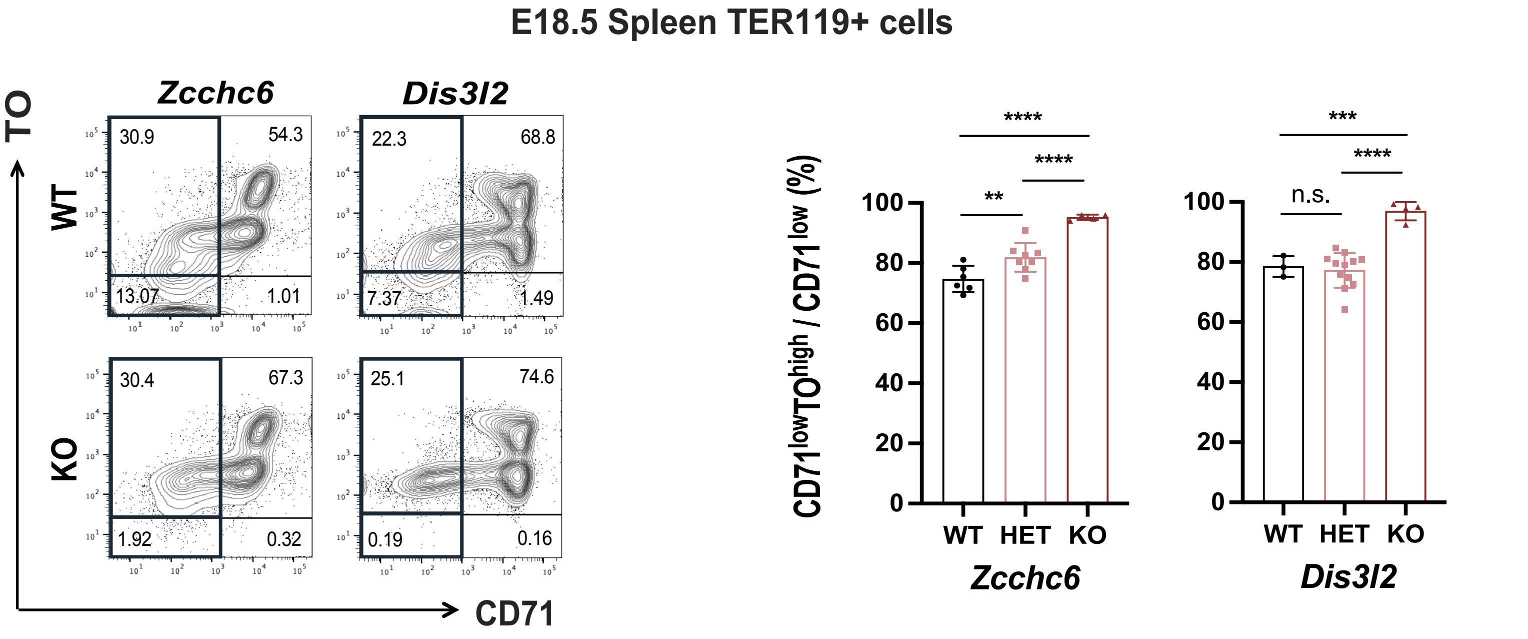


**b**

p-values ( <0.05 *, <0.01 **, <0.001***)

| Parameter | Unit | *Zcchc6* | | | p-value | | *Dis3l2* | | | p-value | |
|---|---|---|---|---|---|---|---|---|---|---|---|
| | | WT | HET | KO | WT vs. KO | HET vs. KO | WT | HET | KO | WT vs. KO | HET vs. KO |
| *Leukocytes:* | | | | | | | | | | | |
| WBC | K/µl | 7.10 ± 1.39 | 8.96 ± 3.72 | 21.13 ± 8.52 | *** | *** | 3.79 ± 1.22 | 4.42 ± 1.08 | 5.53 ± 1.36 | * | |
| LY | K/µl | 5.85 ± 1.25 | 7.60 ± 3.50 | 18.61 ± 7.52 | *** | *** | 1.65 ± 0.62 | 1.96 ± 0.49 | 2.77 ± 1.10 | * | * |
| NE | K/µl | 0.96 ± 0.14 | 0.98 ± 0.31 | 1.66 ± 0.74 | ** | ** | 1.58 ± 0.57 | 1.82 ± 0.57 | 2.00 ± 0.73 | | |
| MO | K/µl | 0.17 ± 0.08 | 0.22 ± 0.07 | 0.71 ± 0.29 | *** | *** | 0.12 ± 0.10 | 0.12 ± 0.09 | 0.15 ± 0.09 | | |
| EO | K/µl | 0.10 ± 0.04 | 0.11 ± 0.05 | 0.12 ± 0.04 | | | 0.34 ± 0.11 | 0.43 ± 0.18 | 0.46 ± 0.10 | | |
| BA | K/µl | 0.04 ± 0.01 | 0.04 ± 0.02 | 0.05 ± 0.03 | | | 0.10 ± 0.05 | 0.10 ± 0.05 | 0.16 ± 0.09 | | * |
| LY | % | 82.10 ± 2.33 | 83.63 ± 5.61 | 87.87 ± 1.61 | | | 43.52 ± 9.67 | 44.52 ± 7.34 | 49.51 ± 11.79 | | |
| NE | % | 13.75 ± 2.22 | 11.93 ± 4.92 | 7.92 ± 1.44 | * | | 41.88 ± 9.86 | 40.86 ± 6.63 | 36.35 ± 9.34 | | |
| MO | % | 2.24 ± 0.92 | 2.60 ± 0.63 | 3.38 ± 0.63 | * | | 2.73 ± 1.84 | 2.75 ± 2.05 | 2.51 ± 1.23 | | |
| EO | % | 1.37 ± 0.44 | 1.37 ± 0.61 | 0.60 ± 0.21 | | * | 9.37 ± 1.14 | 9.64 ± 3.13 | 8.58 ± 2.26 | | |
| BA | % | 0.54 ± 0.06 | 0.47 ± 0.19 | 0.22 ± 0.09 | ** | ** | 2.51 ± 1.15 | 2.23 ± 1.16 | 3.05 ± 1.26 | | |
| *Thrombocytes:* | | | | | | | | | | | |
| PLT | K/µl | 496.75 ± 49.31 | 517.67 ± 35.27 | 519.13 ± 24.93 | | | 425.60 ± 71.20 | 464.71 ± 86.20 | 411.64 ± 66.27 | | |
| MPV | fL | 6.31 ± 0.17 | 6.42 ± 0.15 | 6.50 ± 0.11 | | | 6.32 ± 0.18 | 6.10 ± 0.27 | 6.13 ± 0.22 | | |

### Extended Data Fig. 5

#### Additional Analysis of Conditional *Zcchc6/Dis3l2* KO.

**a**

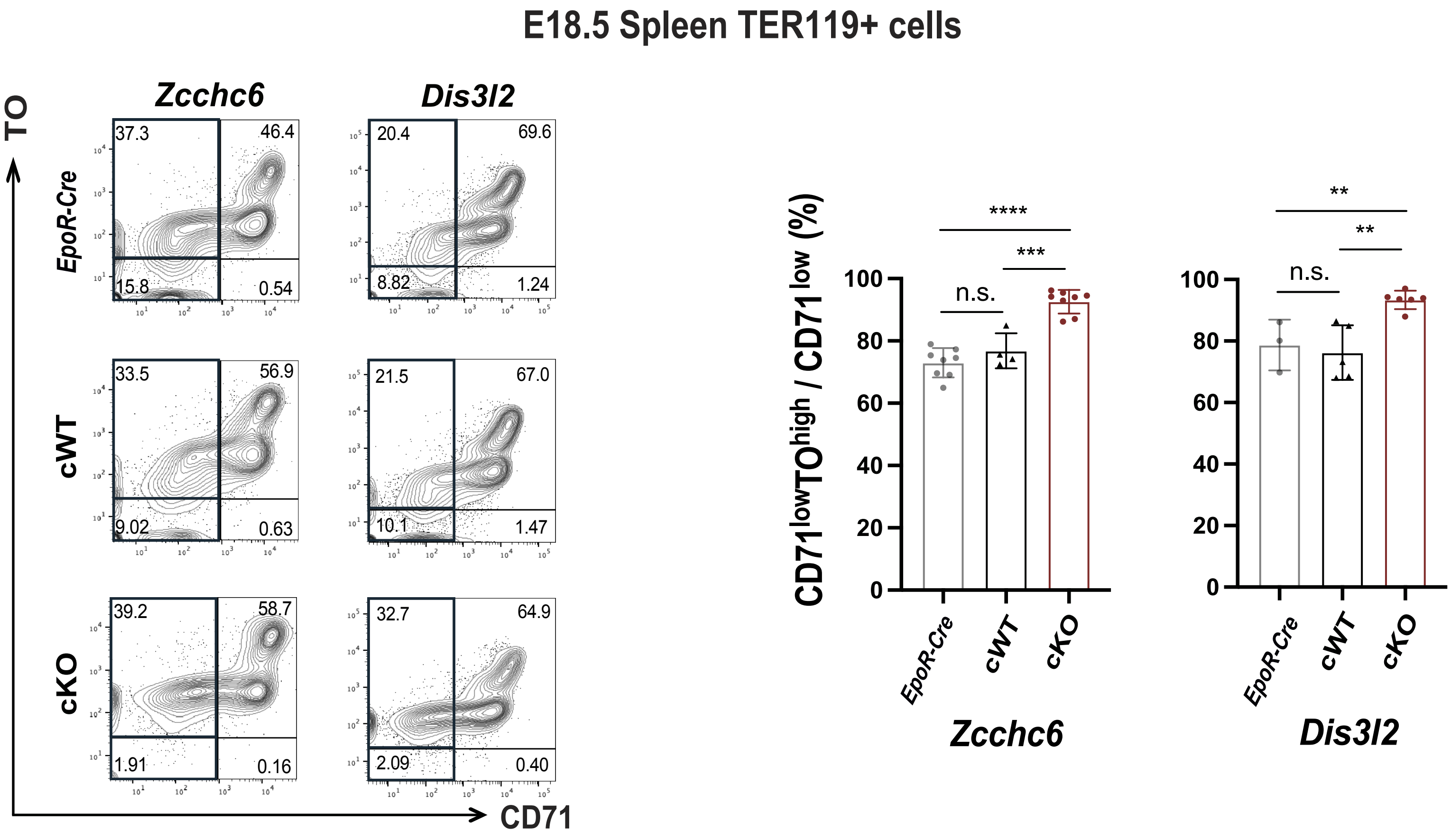


**b**

p-values ( <0.05 *, <0.01 **, <0.001***)

| Parameter | Unit | *Zcchc6* | | | p-value | | *Dis3l2* | | | p-value | |
|---|---|---|---|---|---|---|---|---|---|---|---|
| | | *EpoR-Cre* | cWT | cKO | cWT vs. cKO | *EpoR-Cre* vs. cKO | *EpoR-Cre* | cWT | cKO | cWT vs. cKO | *EpoR-Cre* vs. cKO |
| *Leukocytes:* | | | | | | | | | | | |
| WBC | K/µl | 15.60 ± 5.12 | 16.54 ± 6.04 | 30.89 ± 11.89 | 0.0686 | * | 17.14 ± 15.10 | 13.37 ± 7.68 | 26.49 ± 16.83 | *** | * |
| LY | K/µl | 14.28 ± 5.13 | 15.24 ± 5.72 | 27.93 ± 11.59 | 0.0913 | * | 15.49 ± 14.10 | 12.25 ± 7.40 | 24.19 ± 15.79 | *** | * |
| NE | K/µl | 0.92 ± 0.21 | 0.86 ± 0.20 | 1.86 ± 0.89 | | | 1.18 ± 0.49 | 0.88 ± 0.25 | 1.49 ± 0.52 | ** | |
| MO | K/µl | 0.32 ± 0.13 | 0.36 ± 0.22 | 0.91 ± 0.43 | | * | 0.40 ± 0.55 | 0.17 ± 0.17 | 0.73 ± 0.66 | ** | |
| EO | K/µl | 0.06 ± 0.03 | 0.06 ± 0.03 | 0.16 ± 0.08 | | | 0.06 ± 0.04 | 0.06 ± 0.03 | 0.04 ± 0.02 | | |
| BA | K/µl | 0.03 ± 0.01 | 0.02 ± 0.01 | 0.03 ± 0.01 | | | 0.02 ± 0.01 | 0.02 ± 0.01 | 0.02 ± 0.01 | | |
| LY | % | 90.50 ± 4.55 | 91.88 ± 1.40 | 89.39 ± 4.33 | | | 87.57 ± 7.37 | 88.59 ± 7.74 | 90.05 ± 3.11 | | |
| NE | % | 6.85 ± 4.25 | 5.58 ± 1.66 | 7.21± 4.31 | | | 10.20 ± 7.96 | 9.72 ± 7.94 | 7.41 ± 3.66 | * | |
| MO | % | 1.99 ± 0.27 | 2.02 ± 0.70 | 2.77 ± 1.08 | | | 1.74 ± 0.93 | 0.94 ± 0.64 | 2.21 ± 1.09 | *** | |
| EO | % | 0.47 ± 0.38 | 0.36 ± 0.20 | 0.51 ± 0.13 | | | 0.38 ± 0.18 | 0.56 ± 0.34 | 0.21 ± 0.10 | * | |
| BA | % | 0.20 ± 0.15 | 0.17 ± 0.11 | 0.11 ± 0.08 | | | 0.12 ± 0.06 | 0.17 ± 0.11 | 0.11 ± 0.11 | | |
| *Thrombocytes:* | | | | | | | | | | | |
| PLT | K/µl | 590.75 ± 38.23 | 579.50 ± 16.09 | 609.67 ± 48.13 | | | 610.5 ± 133.57 | 609.11 ± 94.52 | 569.50 ± 57.43 | | |
| MPV | fL | 6.00 ± 0.14 | 6.10 ± 0.18 | 6.02 ± 0.13 | | | 5.98 ± 0.15 | 6.12 ± 0.16 | 6.06 ± 0.28 | | |

### Extended Data Fig. 6

#### Analysis of Hemolytic Anemia Markers.

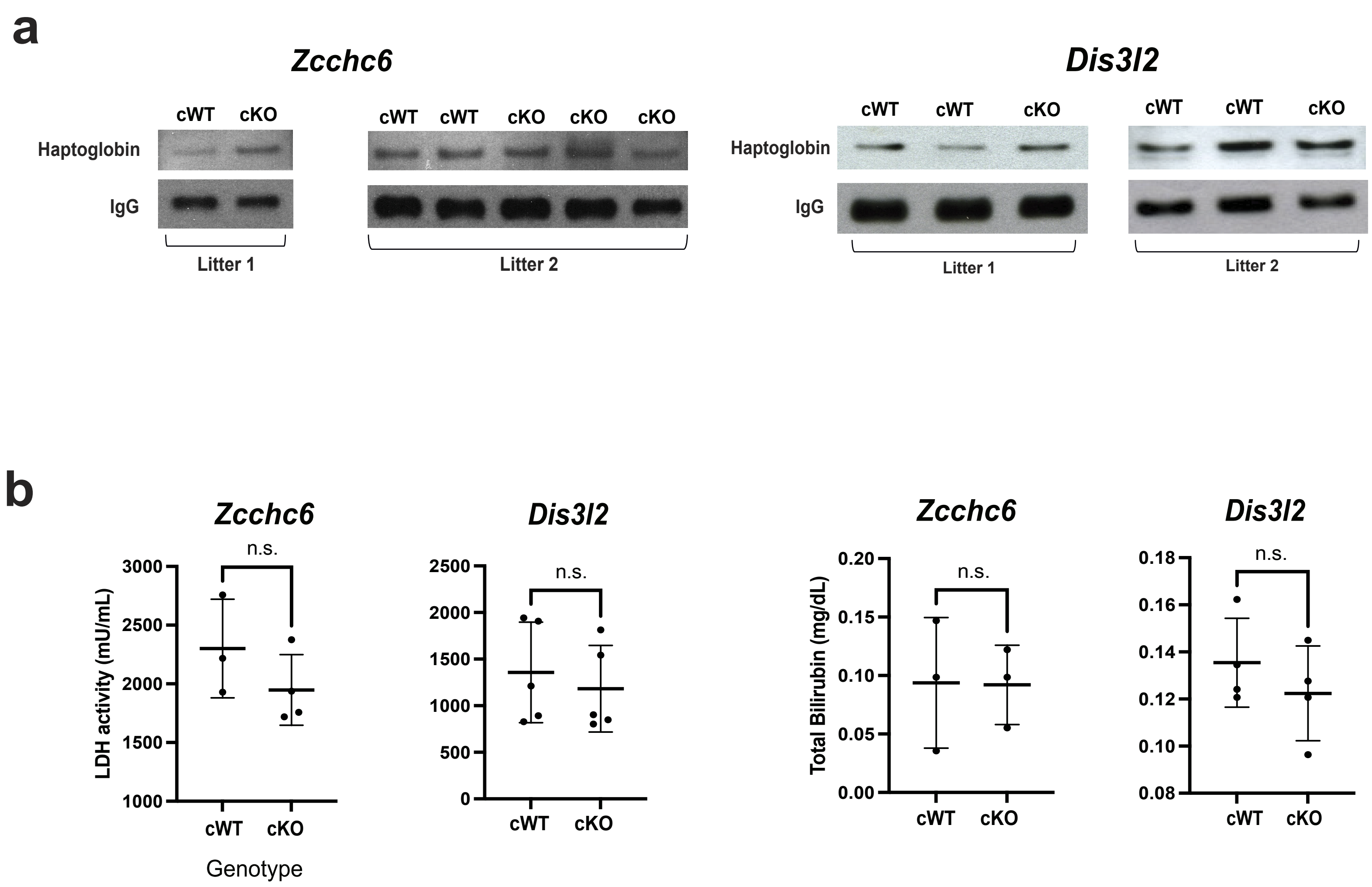

### Extended Data Fig. 7

#### Characterizing the Impact of Knockouts in Pathway-Related Genes.

**a**

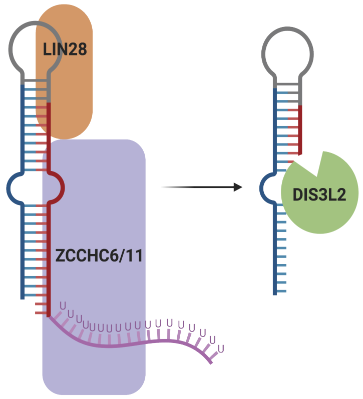


**b**

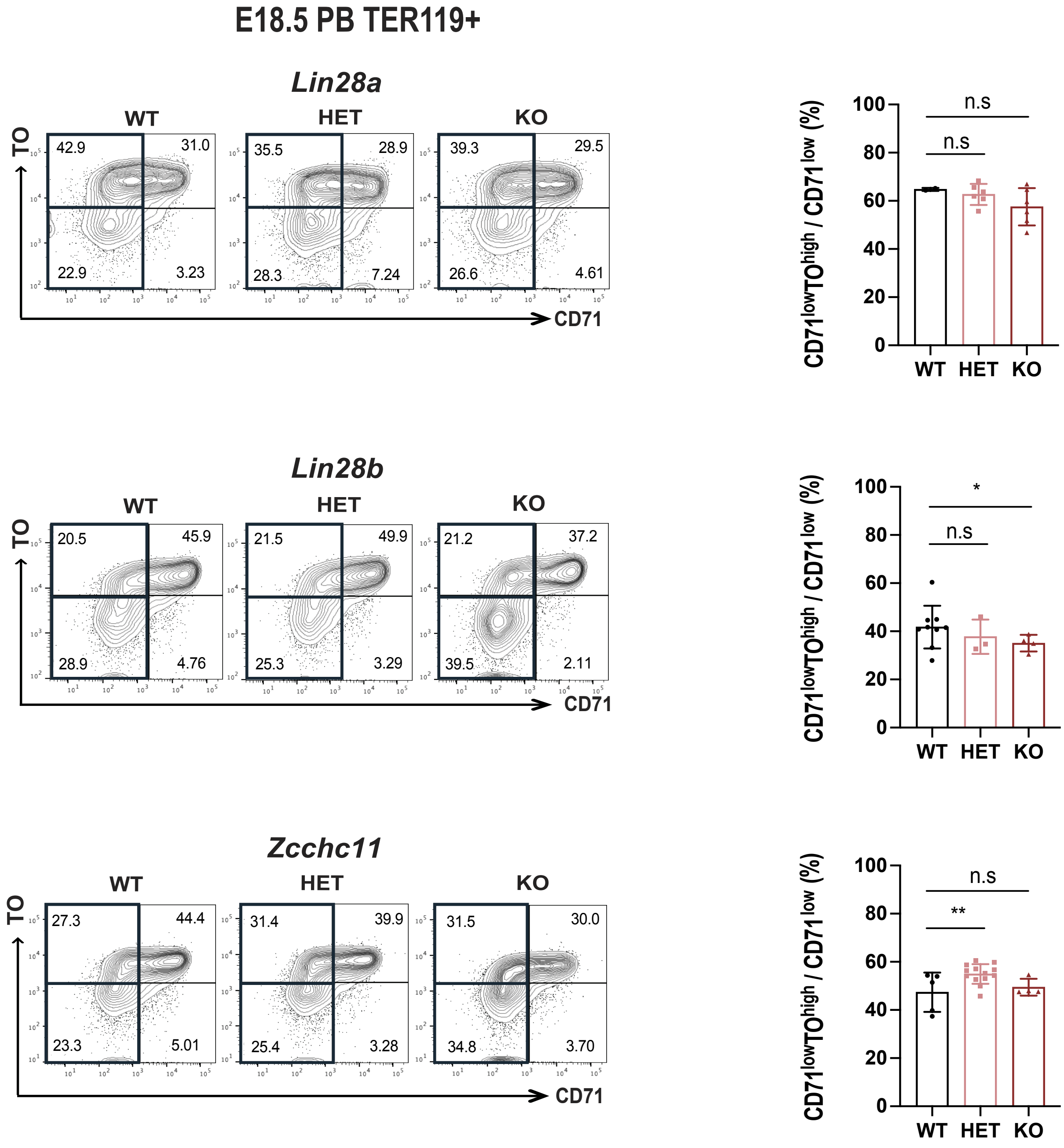

### Extended Data Fig. 8

#### Analysis of Quad-floxed Conditional Knockouts.

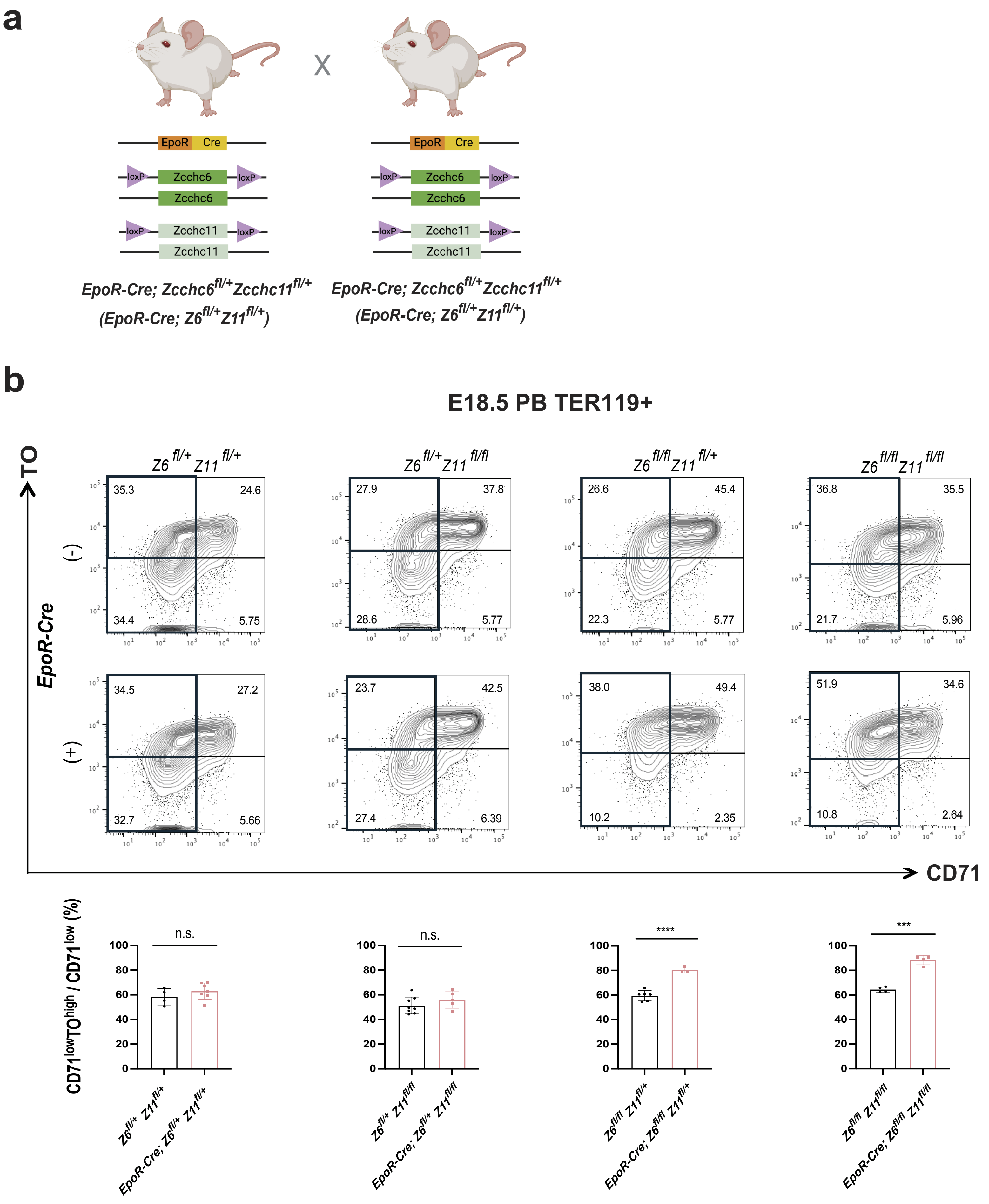

### Extended Data Fig. 9

#### Postnatal Profiling of *Zcchc6* KO and *Dis3l2* cKO.

**a**

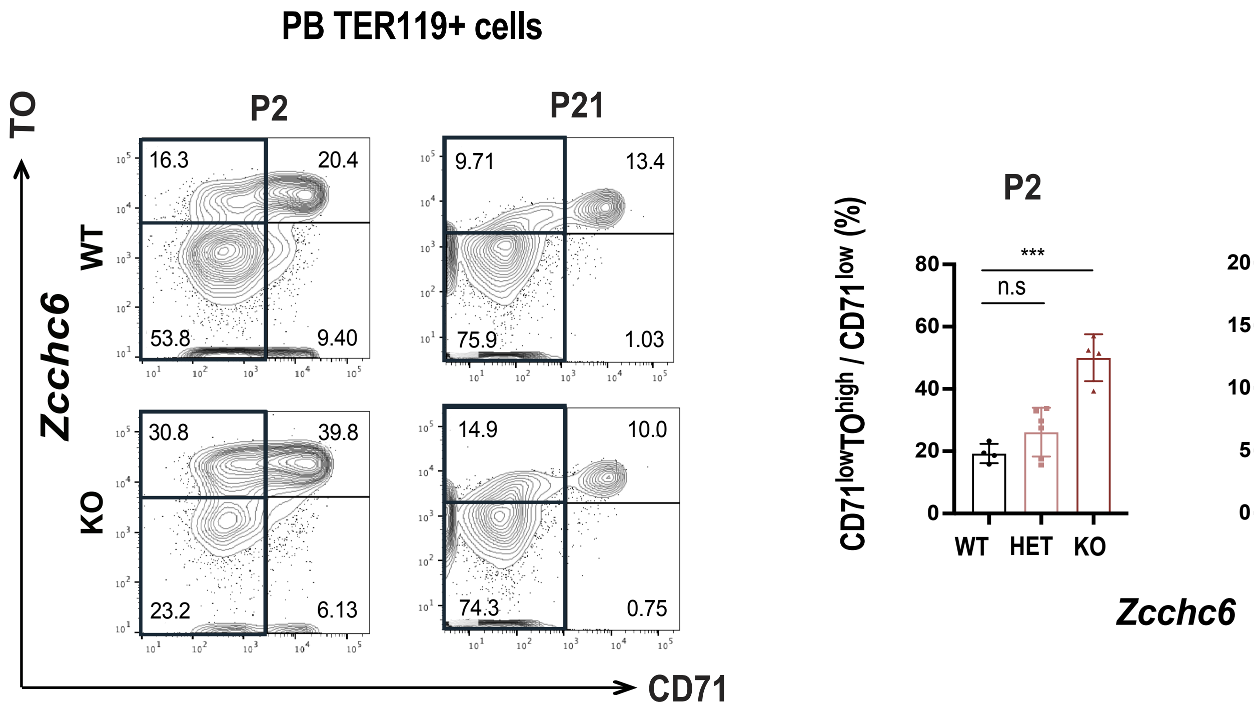


**b**

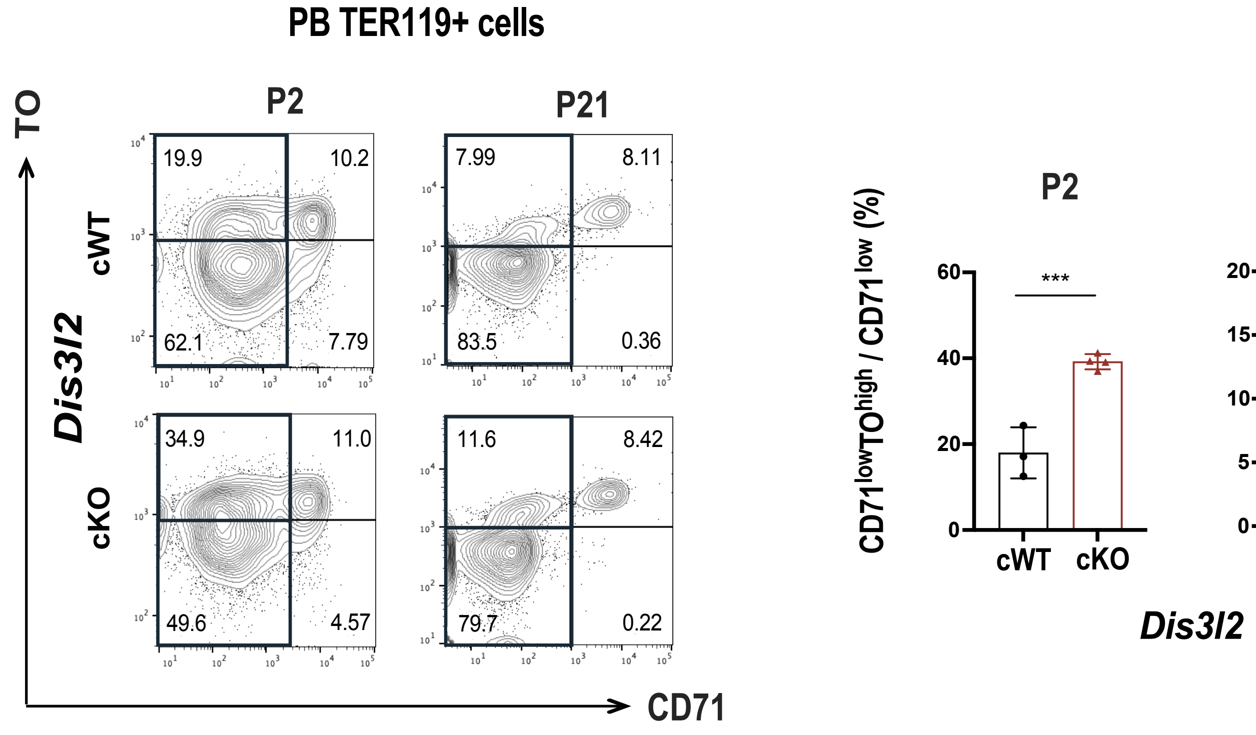

### Extended Data Fig. 10

#### RNA-seq Data Analysis.

**a**

| Genotype | Litter | Total Read No. | Mapped Read No. | Mapping Rate |
|---|---|---|---|---|
| *Zcchc6 WT* | L3 | 78,543,026 | 36,284,397 | 46.2% |
| *Zcchc6 WT* | L4 | 90,445,954 | 51,073,608 | 56.5% |
| *Zcchc6 WT* | L5 | 59,412,646 | 30,752,512 | 51.8% |
| *Zcchc6 KO* | L3 | 81,304,604 | 37,445,722 | 46.1% |
| *Zcchc6 KO* | L4 | 79,495,636 | 44,989,077 | 56.6% |
| *Zcchc6 KO* | L5 | 74,278,610 | 37,835,979 | 50.9% |
| *Dis3l2 WT* | L1 | 68,767,710 | 36,391,345 | 52.9% |
| *Dis3l2 WT* | L2 | 72,958,170 | 35,684,609 | 48.9% |
| *Dis3l2 WT* | L2 | 64,899,346 | 32,206,562 | 49.6% |
| *Dis3l2 KO* | L1 | 76,216,868 | 36,395,571 | 47.8% |
| *Dis3l2 KO* | L2 | 70,791,404 | 27,438,581 | 38.8% |
| *Dis3l2 KO* | L2 | 75,760,250 | 35,673,350 | 47.1% |

**b**

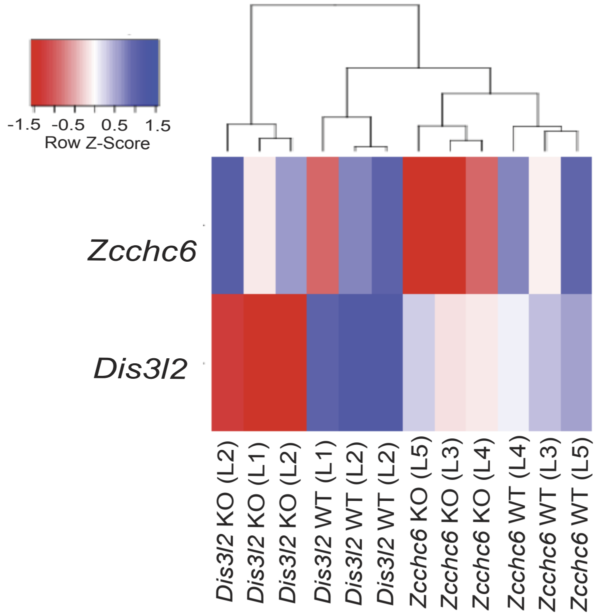


**c**

Enriched Gene Ontology (Biological Process)

Up-regulated upon *Zcchc6* KO

Up-regulated upon *Dis3l2* KO

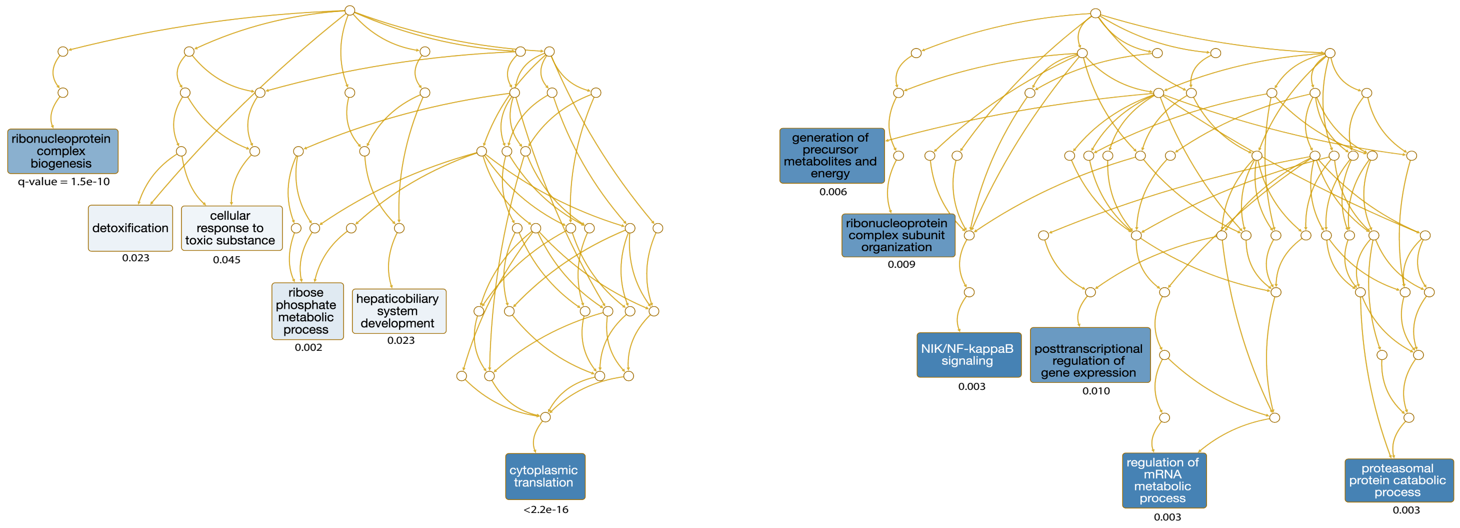

### Extended Data Fig. 11

#### Analysis of Humanized Mice Upon *Zcchc6/Dis3l2* KO.

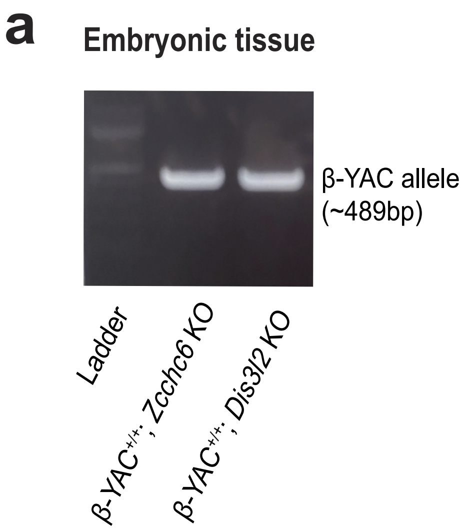


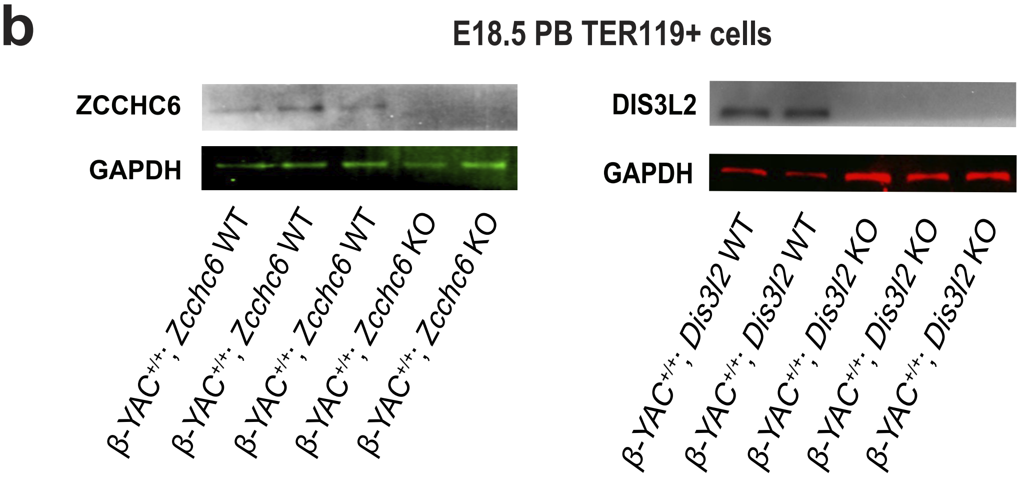


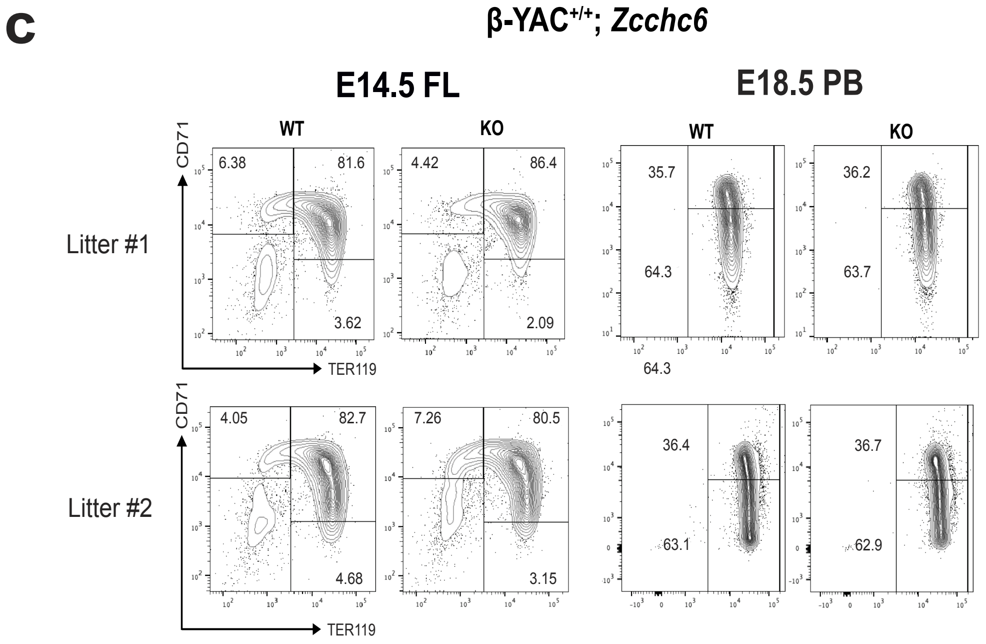


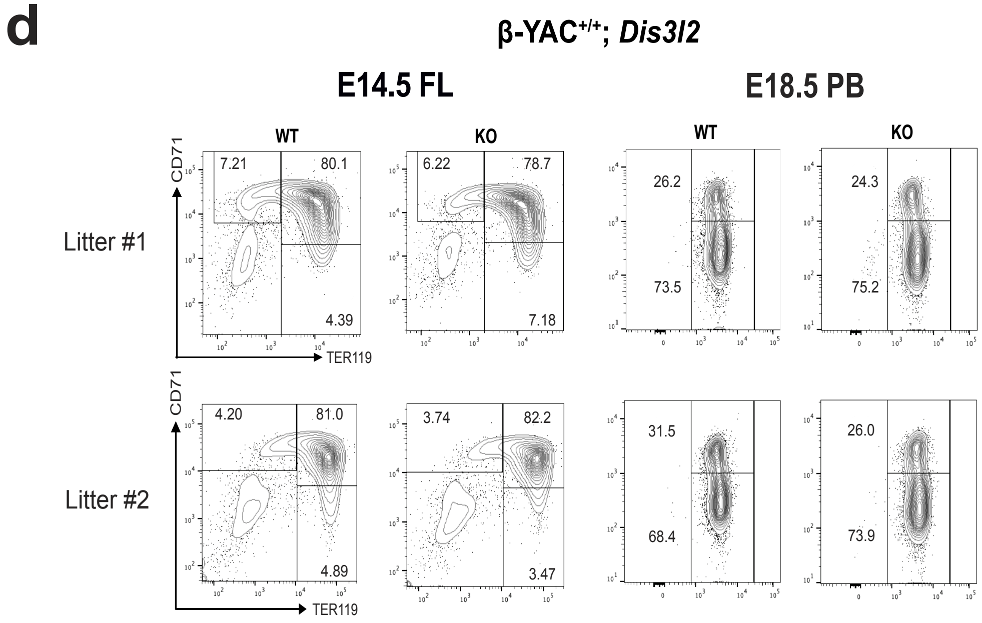

### Extended Data Fig. 12

#### Human *In Vitro* Erythropoiesis

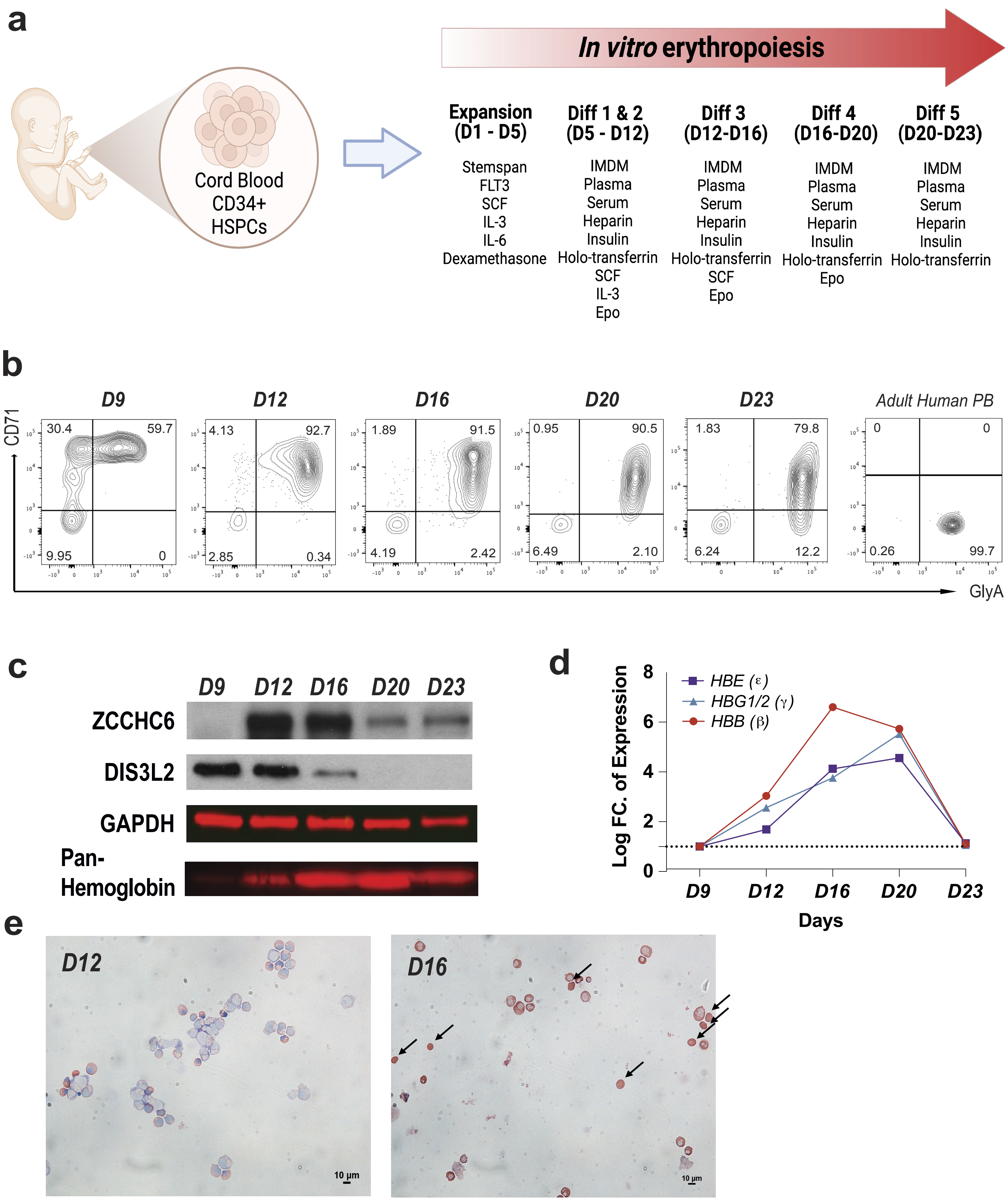

### Extended Data Fig. 13

#### Analysis of CRISPR KO in Differentiated Erythroid Cells from CB.

**a**

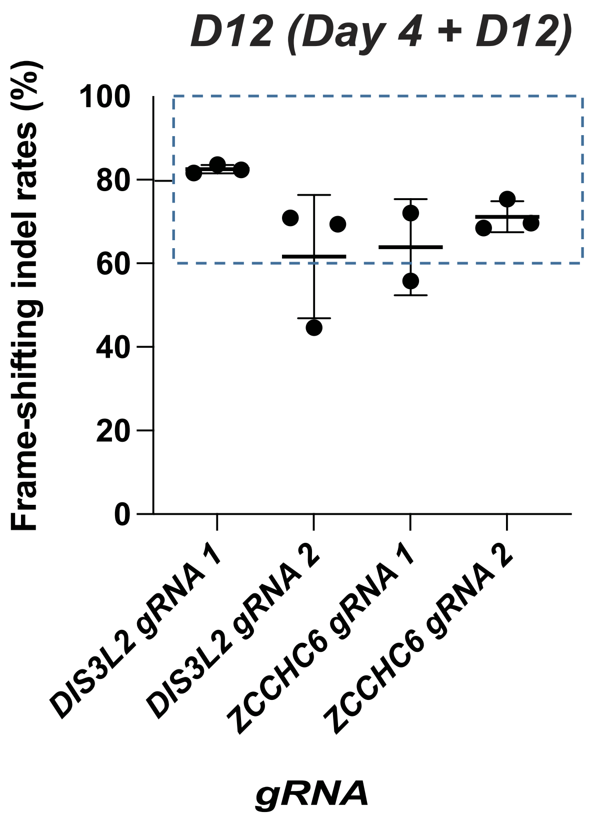


**b**

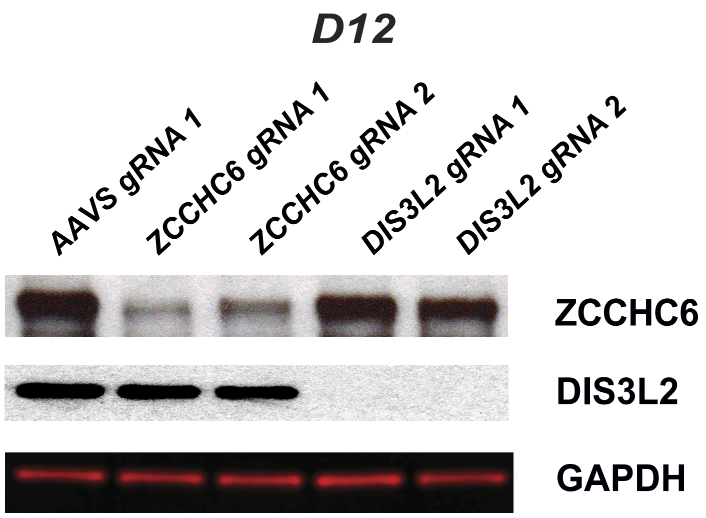


**c**

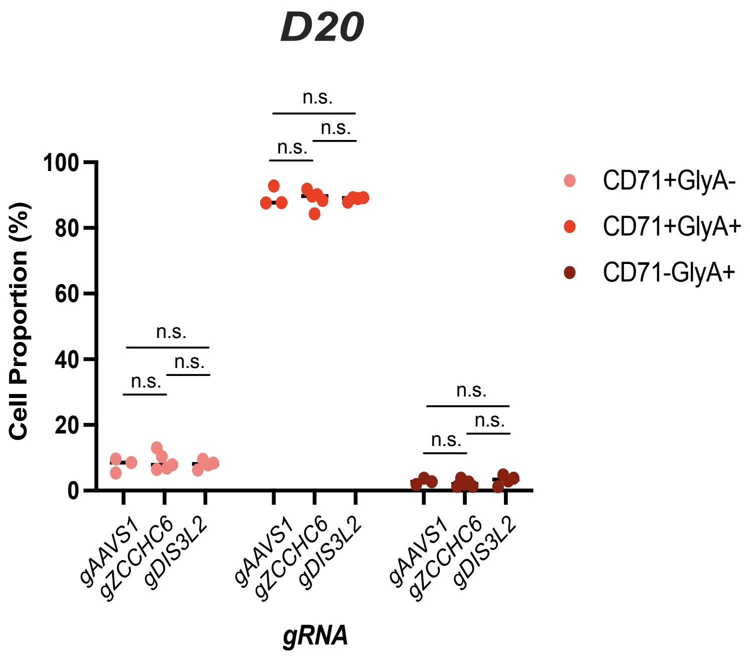

### Extended Data Fig. 14

#### Analysis of CRISPR KO in Differentiated Erythroid Cells from PB Following HU treatment.

**a**

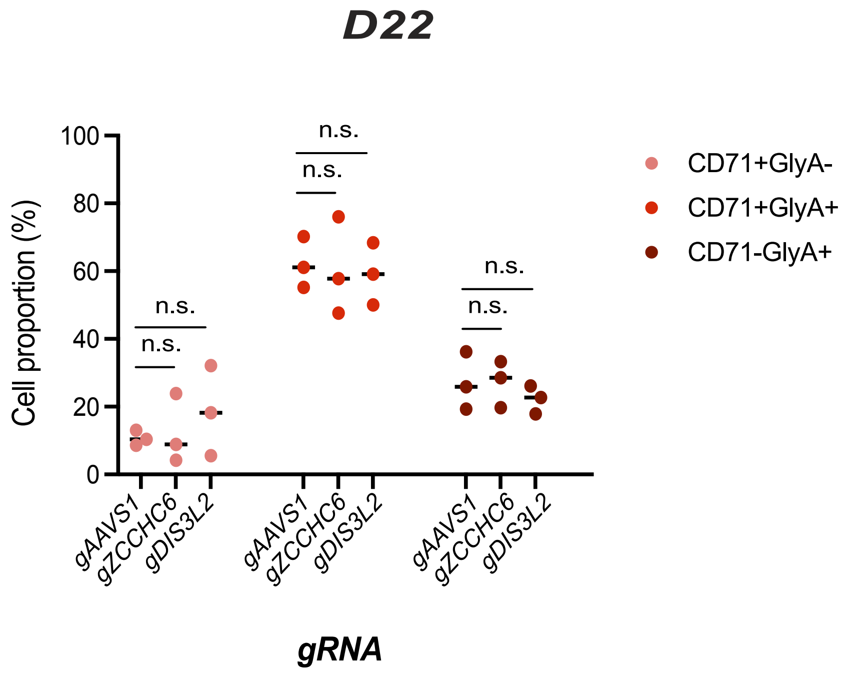


**b**

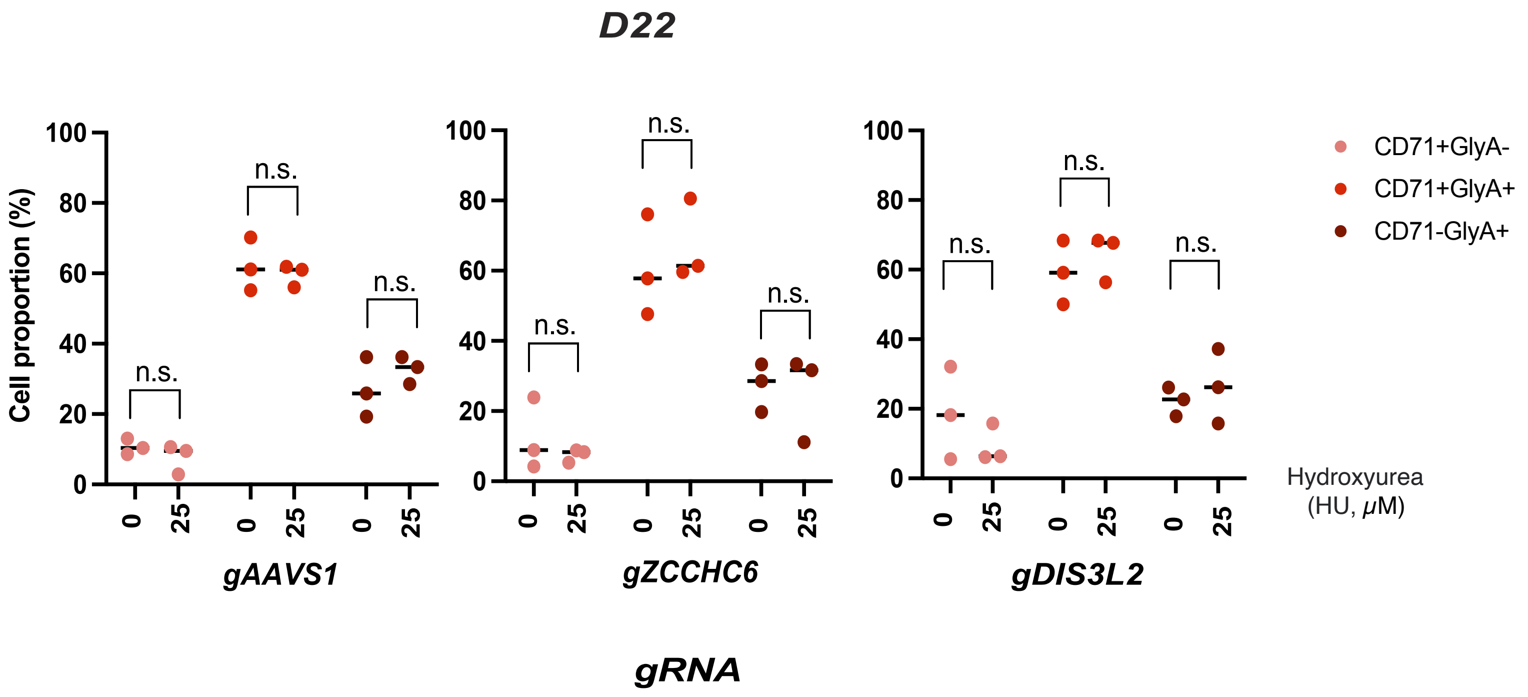
